# Enduring and sex-specific changes in hippocampal gene expression after a subchronic immune challenge

**DOI:** 10.1101/566570

**Authors:** Daria Tchessalova, Natalie C. Tronson

## Abstract

Major illnesses, including heart attack and sepsis, can cause cognitive impairments, depression, and progressive memory decline that persist long after recovery from the original illness. In rodent models of sepsis or subchronic immune challenge, memory deficits also persist for weeks or months, even in the absence of ongoing neuroimmune activation. This raises the question of what mechanisms in the brain mediate such persistent changes in neural function. Here, we used RNA-sequencing as a large-scale, unbiased approach to identify changes in hippocampal gene expression long after a subchronic immune challenge previously established to cause persistent memory impairments in both males and females. We observed enduring dysregulation of gene expression three months after the end of a subchronic immune challenge, Surprisingly, we also found striking sex differences in both the magnitude of changes and the specific genes and pathways altered, where males showed persistent changes in both immune- and plasticity-related genes three months after immune challenge, whereas females showed few such changes. In contrast, females showed striking differential gene expression in response to a subsequent immune challenge. Thus, immune activation has enduring and sex-specific consequences for hippocampal gene expression and the transcriptional response to subsequent stimuli. Together with findings of long-lasting memory impairments after immune challenge, these data suggest that illnesses can cause enduring vulnerability to, cognitive decline, affective disorders, and memory impairments *via* dysregulation of transcriptional processes in the brain.

## 1. INTRODUCTION

The neuroimmune system, and its regulation by peripheral immune states, plays an important regulatory role for behavioral, affective, and cognitive functions (Raison et al., 2006; Tchessalova et al., 2018). In addition to their central role in behavioral and physiological changes during illness and injury (Dantzer et al., 1998), neuroimmune cells and cytokines are activated as a consequence of physical and psychological stressors (Bekhbat and Neigh, 2017), and are critical for normal neural functions including synaptic plasticity and memory (Avital et al., 2003; Yirmiya and Goshen, 2011; del Rey et al., 2013; Adamsky et al., 2018).

Activation of the neuroimmune system as a consequence of surgery, injury, or illness also has long-lasting consequences for memory, affective behaviors, and cognition. In rodent models of sepsis, persistent changes in memory and affective processes have been observed after cecal ligation and puncture (CLP) (Chavan et al., 2012; Huerta et al., 2016) and after high-dose lipopolysaccharide (LPS) treatment (Weberpals et al., 2009; Anderson et al., 2015). More recently, we have demonstrated that a lower-dose subchronic systemic immune challenge results in memory deficits that emerge and persist at least three months after the end of the immune challenge (Tchessalova and Tronson, 2019). These long-lasting memory deficits are likely mediated by changes in neuroimmune and neuronal function that persist long after immune activation and likely contribute to cognitive decline, affective dysregulation, and increased risk of dementia and Alzheimer’s disease long after recovery from major illnesses including sepsis and heart attack (Semmler et al., 2007; Gharacholou et al., 2011; Marra et al., 2018).

A key question is how a transient illness or immune challenge can result in long-lasting changes in memory deficits. One possibility is that neuroimmune signaling persists long after a peripheral immune challenge. Indeed, some sepsis models observe persistent microglial activation and neuroimmune signaling correlate with memory impairments at least one month after surgery or LPS injection (Weberpals et al., 2009; Fenn et al., 2014; Anderson et al., 2015). In contrast, other studies show memory deficits in the absence of ongoing neuroimmune activation in both CLP (Huerta et al., 2016; Singer et al., 2016) and LPS models (Ming et al., 2015; Tchessalova and Tronson, 2019). Here, structural changes in the brain after transient immune challenge, including dysregulation of dendritic spine turnover or density and neuronal loss have been implicated as potential mechanisms for persistent memory impairments (Semmler et al., 2007; Liu et al., 2008; Kondo et al., 2011; Volpe et al., 2015). Additionally, recent work has demonstrated changes in function of seemingly quiescent microglia months after prior immune challenge (Wendeln et al., 2018). Such “priming” (Norden et al., 2015) or “training” of the innate immune system (Netea and van der Meer, 2017) after prior immune experience may result in altered responsivity to both subsequent immune challenge and to other environmental experiences, including learning, stress, or drug exposure.

Yet, if ongoing neuroimmune signaling is not necessary for persistent changes in memory, then how are such long-lasting changes in synaptic function, neuroimmune function, and memory processes maintained long after the immune challenge? One possibility is that these persistent changes in memory are mediated and maintained by enduring alterations of gene expression and associated transcriptional regulatory mechanisms (Tchessalova et al., 2018). Changes in transcriptional regulation as a result of chromatin and DNA modifications (Cholewa-Waclaw et al., 2016) occur after a variety of environmental exposures and life experience, including stress and exposure to addictive drugs, and mediate persistent changes in cellular function and behavior (Gray et al., 2014; Nestler, 2014; Tafet and Nemeroff, 2016).

Here we examined whether changes in hippocampal gene expression persist long after an immune challenge in males and in females and discuss the implications for persistent changes in neural function and dysregulation of memory, emotion, and cognition. We used an unbiased approach, RNA-sequencing, to identify both immune- and non-immune-related gene expression changes long after a systemic immune challenge in both males and females. Since changes in gene expression after experiences such as stress (Gray et al., 2014) are often more evident in response to a subsequent stressor, we examined changes in gene expression both months after a subchronic immune challenge, and in response to a subsequent immune challenge. We found that a transient, subchronic LPS protocol (Tchessalova and Tronson, 2019) resulted in persistent and sex-specific dysregulation of gene expression in the hippocampus. Males and females differed in both number and patterns of gene expression, where males show more changes in baseline gene expression and females show greater changes in response to an additional, acute challenge. These data have important implications for identifying the precise mechanisms by which a prior illness or injury can impact on cognitive function and vulnerability to cognitive decline and dementia.

## 2. MATERIALS AND METHODS

### 2.1 Animals

Male and female 8-9 week old C57BL/6N mice were purchased from Envigo (Indianapolis, IN). All mice were individually housed with *ad libitum* access to standard mouse chow and water as individual housing in mice prevents fighting-induced stress in males (Meakin *et al*, 2013) and is ethologically appropriate for males and females (Becker and Koob, 2016). The facility is ventilated with constant air exchange (60 m^3^/ h), temperature (22 ±1°C), and humidity (55±10%) with a standard 12 h light-dark cycle (Keiser *et al*, 2017). All animals were previously tested on context fear conditioning (Tchessalova and Tronson, 2019), anxiety-like behavior, and forced swim test. We waited one month after behavioral testing to avoid assessing interactions of acute behavioral tasks with persistent effects of prior immune challenge. All experimental methods used in these studies were approved by the University of Michigan Committee on the Use and Care of Animals. Sample size was n = 6 per immune challenge condition, with 3 males and 3 females per group.

### 2.2 Subchronic Immune Challenge

Lipopolysaccharide (LPS, *Escherichia coli*, serotype 0111:B4; Sigma-Aldrich, St. Louis) was dissolved in saline for a final concentration of 12.5μg/mL. LPS injections were administered intraperitoneally (i.p.) at a dose of 250μg/kg. Vehicle control mice received an equivalent volume of saline (20mL/kg, i.p.) (Cloutier et al., 2012; Tchessalova and Tronson, 2019). The subchronic immune challenge consisted of five injections, spaced three days apart (days 1, 4, 7, 10, 13). All injections were administered in the morning between 9 and 10am. In the Long-Term condition, tissue collection occurred 12 weeks after the final LPS injection. Separate animals received a subsequent LPS (250μg/kg; or vehicle) injection 12 weeks after the subchronic immune challenge and tissue was collected 6 hours after injection (Long-term + Acute condition).

### 2.3 RNA extraction and sequencing

To maintain RNA quality, whole hippocampi were collected and immediately placed in RNALater until processing (Hodes et al., 2015; Bagot et al., 2016; Lorsch et al., 2018; Walker et al., 2018). One hemisphere (counterbalanced by side between sex and experimental conditions) was selected for RNA extraction while the other was collected for protein analysis. RNA was isolated using Life Technologies PureLink RNA Mini kit (cat. no. 12183018A). Relative RNA quantity and integrity were first analyzed using NanoDrop (ThermoFisher) and gel electrophoresis. Quality and integrity checks were then completed on the Bioanalyzer by the University of Michigan DNA sequencing core, with acceptable RIN values greater than 7. Sequencing was performed using Illumina 4000 High-Seq platform, using single-end, nonstrand, Ribo-depletion with read lengths of 50 and sequencing depth of 40 million reads per sample. Total RNA (20ug) was used to construct the mRNA libraries. Barcoded cDNA libraries were constructed from polyadenylated transcripts that were purified, fragmented, and reverse transcribed using random hexameters.

### 2.4 Differential gene expression analyses

Alignment, differential expression analysis, and post-analysis diagnostics were analyzed using the Tuxedo Suite software package. Reads were aligned to the Ensembl *Mus musculus* NCBIM37 reference genome using TopHat and Bowtie. The quality of the raw reads data for each sample was assessed using FastQC to exclude any reads with quality problems. Expression quantitation, normalization, and differential expression analyses were conducted through Cufflinks/CuffDiff with UCSC mm10.fa reference genome sequence. Differentially expressed genes (DEGs) were determined by multiple comparison correction using FDR > 0.05 cutoff.

#### Visualization differentially expression genes

DEGs in males and females were visualized using Venn diagrams through Venny 2.1.0 (http://bioinfogp.cnb.csic.es/tools/venny/). Volcano plots generated through Advaita’s iPathway guide (https://www.advaitabio.com/ipathwayguide) were used to view DEGs by fold change and significance (q-value).

#### Biological pathways

Metascape (http://metascape.org/gp/index.html#/main/step1) was used to generate gene annotation and gene list enrichment analysis, with a focus on biological pathways and processes. Significance of biological processes was determined through P-values calculated on a hypergeometric distribution (log10). Reference gene lists and annotated information were obtained from the Enrichr web page. Metascape was also used to conduct meta-analysis, with generation of Circos plots to visualize shared genes and pathways amongst the conditions and heatmaps of gene ontology (GO) terms that hierarchically cluster together amongst experimental conditions.

#### Protein-protein interactions

Clusters of functional protein-protein interactions (existing and predicted) between targets in experimental conditions of interest were visualized using the STRING 10.5 software (https://string-db.org/).

#### Statistical analysis

Criteria for a differentially expressed gene included a fold change greater than 1.5, and false discovery rate greater than 5% (fold change ≥ ± 1.5 and FDR ≤ 0.05). All comparisons of DEGs between control and experimental conditions were made within sex using unpaired *t* tests (two-tailed) with Benjamini correction to account for multiple comparisons to determine the effect of immune challenge in males and in females separately. DEGs in the hippocampus of males and females at baseline were compared using unpaired *t* tests (two-tailed) between vehicle treated males and vehicle treated females. Groups were compared as shown in Table 1.

**Table 1.**
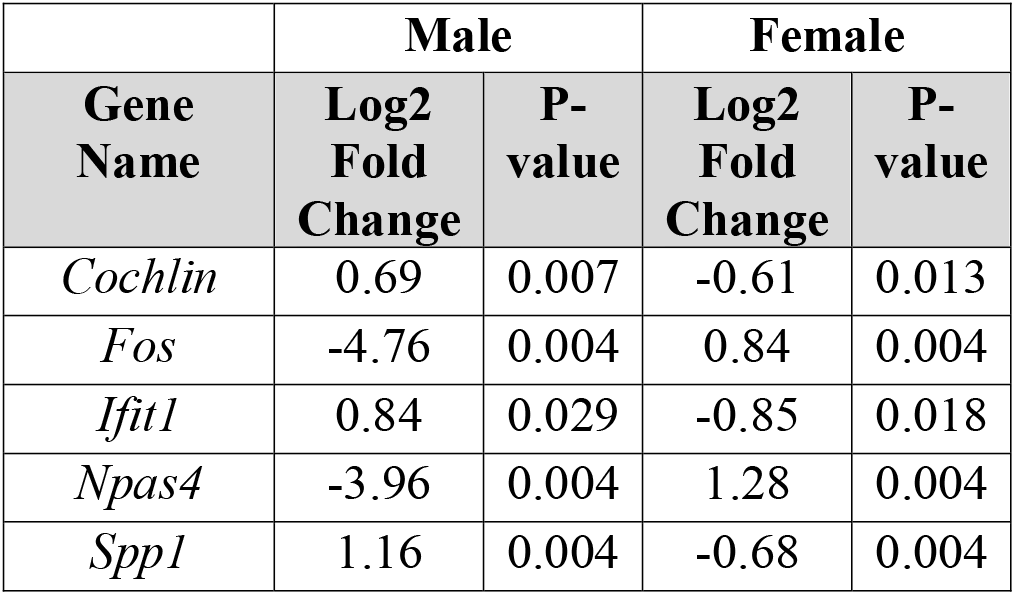
Differentially expressed genes in the hippocampus of males three months after subchronic immune challenge (Long-term condition).

#### Data availability

To allow all interested parties to explore and utilize our processed data, we have made our data publicly available through user-friendly databases, including the Gene Expression Omnibus (GEO), with accession number GSE126678, and Sequence Read Archive (SRA), with SRA number SRP186132 and BioProject number PRJNA522922.

## 3 RESULTS

### 3.1 Subchronic immune challenge induces persistent changes in gene expression

We observed striking changes in gene expression in the hippocampus 12 weeks after subchronic immune challenge in males, with fewer changes observed in females. Of over 20,000 genes detected, there were 230 DEGs in the hippocampus of males and 26 DEGs in the hippocampus of females (Figure 1A). In males, 183 genes were significantly upregulated and 47 significantly downregulated (see Supplementary Table 1 for a full list of DEGs). In females, 7 genes were significantly upregulated and 18 downregulated (see Supplementary Table 2 for a full list of DEGs). Five of these genes were differentially expressed in both males and females, with *Npas4* and *fos* downregulated in males and upregulated in females, and *Ifit1, Spp1* (*Opn*), and *Coch* upregulated in males and downregulated in females (Figure 1; Table 1). Overall, in males, neuroimmune-related genes showed persistent upregulation, whereas neuroplasticity-related genes showed downregulation months after the subchronic immune challenge. The 10 most upregulated genes, included *Wdr72, Prdm6, Slc16a8, Tmem72, Wfdc2, Cldn2, Kcne2, Steap1, Ttr*, and *Aqp1* while the 10 most downregulated genes included immediate-early genes and transcription factors *Egr2, Fosb, Fos, Npas4, Egr4*, and *Junb* as well as genes related to immune signaling and transcription, *Btg2* and *Ccl3*, and extracellular matrix associated *Cyr6* (Figure 1B). Other upregulated genes of interest, with lower log2 fold change values included plasticity-related *Cdh3*, tight-junction proteins such as *Cldn9*, and targets related to G-protein signaling (e.g. *Ccdc135*). Additional downregulated genes included other neuroplasticity related genes, such as *Nr4a1* and *Dusp6*, and cytokine *Ccl3* (Figure 1C) as well as immune-related *Ier2* and *Apold1*.

**Figure 1.**
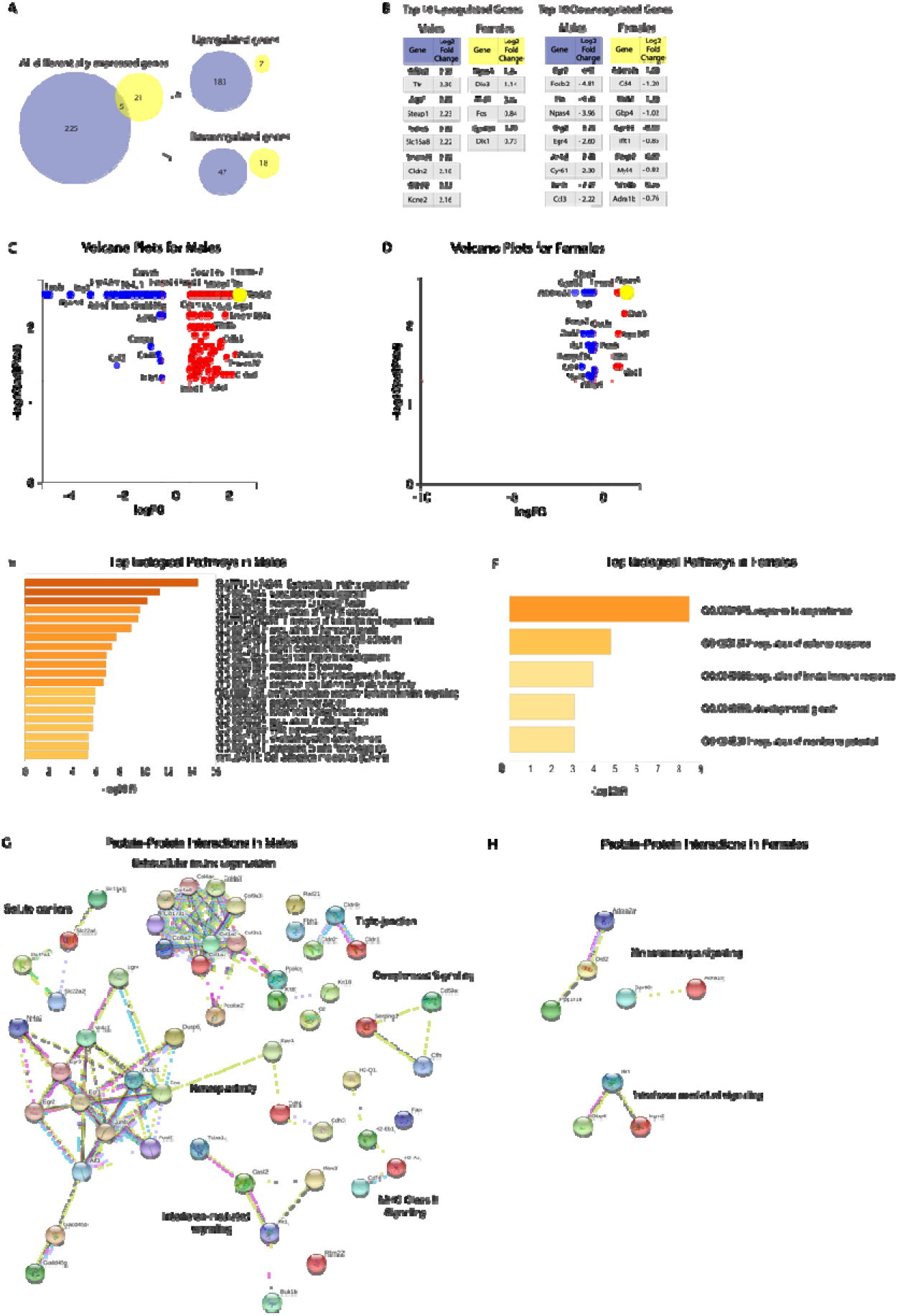
Subchronic, peripheral LPS challenge induces changes in hippocampal gene expression 12 weeks after last injection. (A) Males (purple) show a greater number of DEGs than females (yellow). Males showed 183 upregulated and 47 downregulated genes, whereas females showed 7 and 18 up- and down-regulated genes, respectively. (B) Top upregulated and downregulated genes in males and females. (C, D) Volcano plots showing differentially expressed genes in males (C) and females (D) obtained from Advaita iPathway Analysis. Differentially expressed genes (DEG) are represented in terms of their measured expression change (x-axis) and the significance of the change (y-axis), with upregulated genes shown in red and downregulated genes shown in blue. (E,F) Biological pathways and processes enriched in gene set (A) in males and (B) in females were generated through Metascape. Distinct biological pathways are observed in males and females, with greater plasticity and immune-related targets in males and greater monoaminergic signaling in females. (G, H) Protein-protein interaction (PPI) networks of targets were generated using STRING 10.5 and clustered by biological function (G) in males and (H) in females. Edges between nodes are color coded for relationship type. Blue: known interactions; Pink experimentally determined interactions; Black: Co-expression of targets; Purple: protein homology; Green, yellow, and dark blue: predicted interactions.

In females, although few DEGs were identified, they were consistently related to dopaminergic/monoaminergic signaling. The top upregulated genes included *Npas4, Dio3, Mid1, Fos, Gpr101, and Dlk1* while the top 10 downregulated genes included immune-related and monoaminergic-associated *Adora2a, Cd4, Drd2, Gbp4, Gpr88, Ifit1, Foxp2, Myl4, Scn4b*, and *Adra1b* (Figure 1). Other dysregulated targets of interest included neuropeptide *Penk* and those involved in metabolic functions (e.g. *Xdh*).

Biological pathway and process enrichment analysis revealed top clusters in males were: extracellular matrix organization (log10(P) = −15.41), vascular development (log10(P) = −13.85), response to growth factor (log10(P) = −11.53), regulation of MAPK signaling (log10(P) = −10.2), and response to hormone (log10(P) = −8.87) (Figure 1E). In females, top biological processes included response to amphetamine (log10(P) = −8.49), regulation of defense response (log10(P) = −4.76), and regulation of membrane potential (log10(P) = −3.06) (Figure 1F). Importantly, DEGs in males and females pertain to distinct biological pathways, with neuroplasticity- and MAPK-associated signaling in males and monoaminergic signaling and innate immune-related signaling in females.

Functional protein-protein interaction analysis (STRING) identified groups of DEGs with functional relationships. In males, these clusters included synaptic plasticity and memory-related genes, extracellular matrix targets including collagens and matrix breaking enzymes, growth factors and their receptors, and immediate early genes. There were also several immune-related targets involved in innate immune responses, including regulators of MAPK signaling, complement genes and their regulators, and MHC II-related targets. Notably, some regulators of immune signaling, including *NR4a1/2, Spp1*, and *Dusp1/6* are also plasticity related genes. Smaller clusters of genes included targets associated with organic anion/cation solute carriers, and potassium and chloride channels involved in inhibitory transmission (Figure 1G). In females, STRING analysis identified two clusters: one including dopaminergic signaling, adenosine signaling, adrenergic signaling, and G protein-coupled signaling; and other genes involved in interferon-mediated signaling (Figure 1H). Importantly, the clusters identified in females did not overlap with those identified in males.

### 3.2 Prior subchronic immune challenge alters hippocampal gene expression in response to a later acute LPS injection in a sex-specific manner

We examined the long-lasting effect of prior subchronic immune challenge on transcriptional regulation to a subsequent acute LPS injection (Long-term+Acute condition). In males, we observed 58 DEGs (see Supplementary Table 3), whereas in females, 432 genes were differentially expressed compared with acute immune challenge in previously naïve mice (see Supplementary Table 4). The top 10 upregulated genes in males included *Isl1, Prdm2, Gpr151, Eomes, Barhl2, Chrna3, Slc10a4, Sstr5, Chrnb3, Sln* and the only two downregulated genes included *Fermt1 and Lcn2* (Figure 2). Other upregulated targets with lower log2 fold change include neurotransmitter-associated *Drd1, Chrnb4*, insulin-related *Irs4*, and transcription factor *Foxp2* while the genes with the lowest log2 fold change include G-protein signaling *Gpr153, Adcyap1*, and TGF-beta associated, DNA-binding *Peg10* (Figure 2C. Only *Lcn2* and *Fermt1* were downregulated.

**Figure 2.**
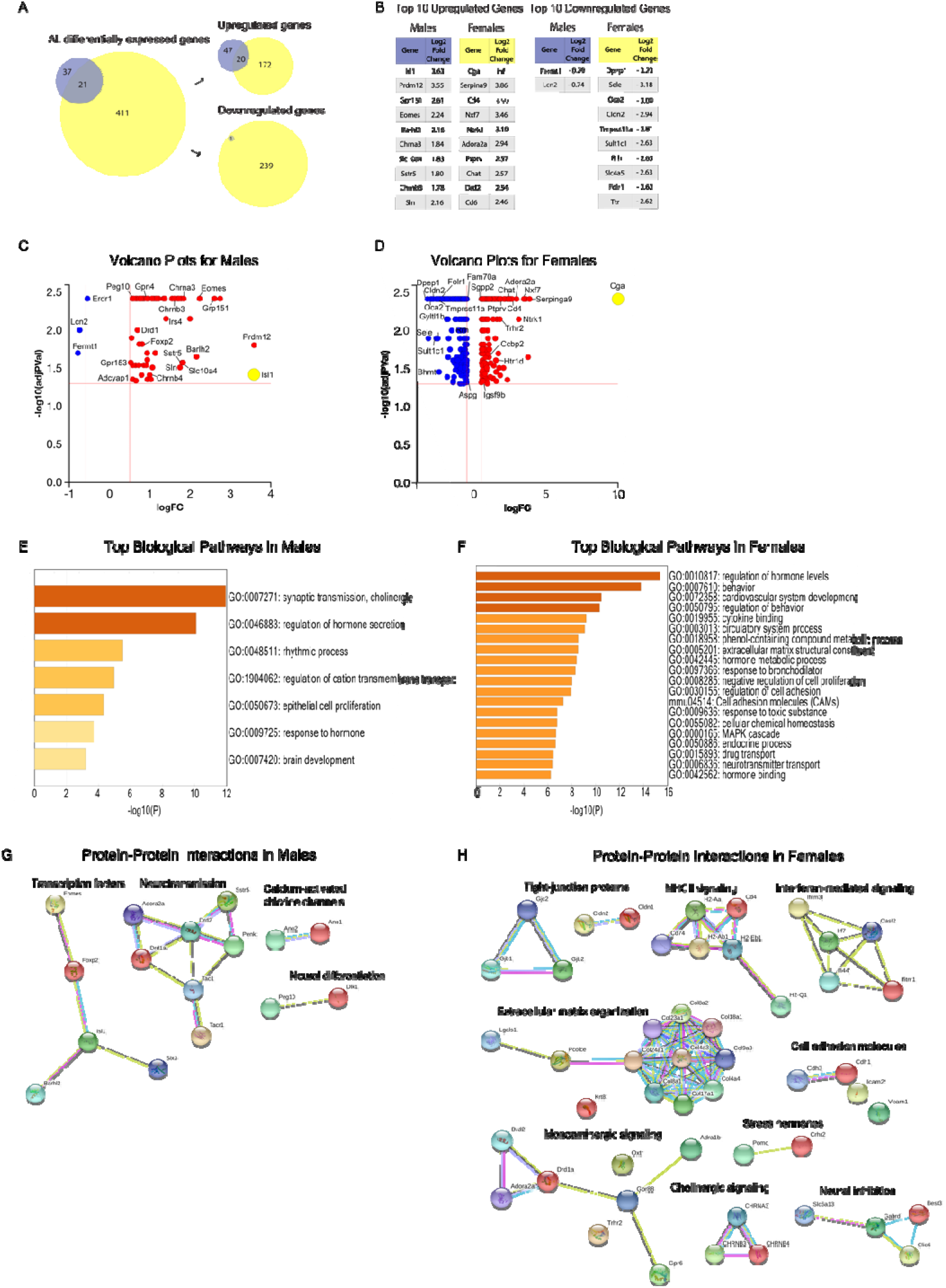
Prior subchronic, peripheral LPS challenge alters hippocampal gene expression in response to a subsequent, acute challenge in a sex-specific manner. (A) Females (yellow) show a greater number of differentially expressed genes than males (purple). Females show 192 upregulated and 240 downregulated genes, whereas males show 67 up- and 1 downregulated gene compared with the response to acute immune challenge in previously naïve animals. (B) Top upregulated and downregulated genes in males and females. (C, D) Volcano plots showing differentially expressed genes in males (C) and females (D) obtained from Advaita iPathway Analysis. Differentially expressed (DE) genes are represented in terms of their measured expression change (x-axis) and the significance of the change (y-axis), with upregulated genes shown in red and downregulated genes shown in blue. (E,F) Distinct biological pathways and processes are enriched in DEGs of (E) males and (F) females who showed greater numbers of pathways and changes in immune-related pathways. (G,H) Protein interaction (PPI) networks of targets, clustered by biological function (G) in males and (H) in females.

In females, the 10 most upregulated genes have immune-related, RNA-binding, and hormone-related functions and included *Cga, Sperina9, Cd4, Nxf7, Ntrk1, Adora2a, Ptprv, Chat, Drd2, Cd6* (Figure 2B). The 10 most downregulated targets were related to extracellular receptor or transporter activity and hormonal or metabolic functions and included *Dpep1, Sele, Oca2, Cldn2, Tmprss11a, Sult1c2, Rrh, Slc4a5, Fol1r*, and *Ttr* (Figure 2B). Additional upregulated targets of interest included neurotransmitter-associated *Htrd1*, hormone-related *Trhr*, immune-associated *Ccbp2*, while other downregulated genes of interest include metabolism-associated *Gytl1b*, and methylation-associated *Bhmt1*, as well as immune-related *Il-31ra* and *Tgfbi*, tight junction associated *Cldn9*, and hormone-associated *Mc3r*. Twenty-one genes, mostly pertaining to neurotransmitter function, showed differential expression in both sexes under these conditions, with 15 genes (e.g. *Gpr151, Chrna/b3, Ptprv, Slc5a7, Tac, Six3, Penk, Drd2, Adora2a, Foxp2, Syt6, Lrrc55, Drd1a, Susd2*) upregulated in both sexes, one gene (*Lcn2*) downregulated in both sexes, and five were dysregulated in opposite directions (e.g. *Slc35a3, AW551984, Arhgap6, Dlk1, Gpx3*) (Figure 2A; Supplementary Table 5).

Biological pathway and process enrichment analysis revealed diverging pathways in males and females (Figure 2E-H). In males, we observed changes in gene expression related to cholinergic synaptic transmission (log10(P) = −11.94), hormone secretion (log10(P) = −10.09), neuropeptide signaling (log10(P) = −7.45), response to nicotine (log10(P) = −6.01), and organic hydroxy compound transport (log10(P) = −6.00) (Figure 2E). In females, the top biological pathways and processes included regulation of hormone levels (log10(P) = −15.39), extracellular matrix organization (log10(P) = −8.1), hormone metabolic processes (log10(P) = −8.41), regulation of cell adhesion (log10(P) = −7.91), and MAPK cascade (log10(P) = −6.68) (Figure 2F).

Functional protein-protein interaction analysis (STRING) in males identified clusters associated with neurotransmission and neuroplasticity, including receptors important for dopaminergic and adenosine receptor-associated signaling and neuropeptides/neuropeptide receptors, a large cluster of transcription factors, and smaller clusters of calcium-activated chloride channels, and neural differentiation (Figure 2G). In females, we found more immune-related clusters than in males, including MHC II signaling, interferon-mediated signaling, cholinergic signaling, and stress hormones. There were also immune and plasticity-related targets, including extracellular matrix and cell adhesion molecules. There were clusters with monoaminergic signaling including dopamine, serotonin, noradrenaline, and adenosine receptors, neuropeptide, and hormone receptors. Smaller clusters included channels, solute carriers important for neuronal inhibition, and metabolic functions (Figure 2H).

### 3.3 Differential gene expression in male versus female hippocampus

We also examined sex differences in baseline gene expression in the hippocampus. We observed 220 genes differentially expressed in the hippocampus of males compared with females, 95 of which are more strongly expressed in males and 125 in females. As expected, Y-chromosome genes including *Ddx3y* were abundant in males but not evident in females, whereas genes that escape from x-inactivation, including *Xist*, were more abundant in females. The top 10 highest genes in males included Y-linked *Ddx3y*, and *Ei2s3y*, and transcription factors *Fos, Fosb, Npas4, Btg2, Egr4*, and others such as *Tmem72, Ttr, Slc4a5;* whereas the top 10 highest genes in females included *Sv2c, Coch, Bmp6, H2-D1, Kdm6a, Nid1, Serpinf1, Tmem90a, Ranbp3l*, and *Plxdc1* which have various functions, including metabolic, immune, and transcriptional roles (Figure 3; Supplementary Table 6). Other genes of interest higher in males included extracellular matrix associated *Krt18* and *Serpine1*, neurotransmitter-associated targets (e.g. *Chrm5*), transcription factors (e.g. *Maff*), and those with metabolic functions (e.g. *Steap1*), while other genes higher in females included MHC class II targets (e.g. *H2-Aa, H2-Q6, Cd4*), interferon signaling-related *ifi47*, and solute carriers (e.g. Slc26a7).

**Figures 3.**
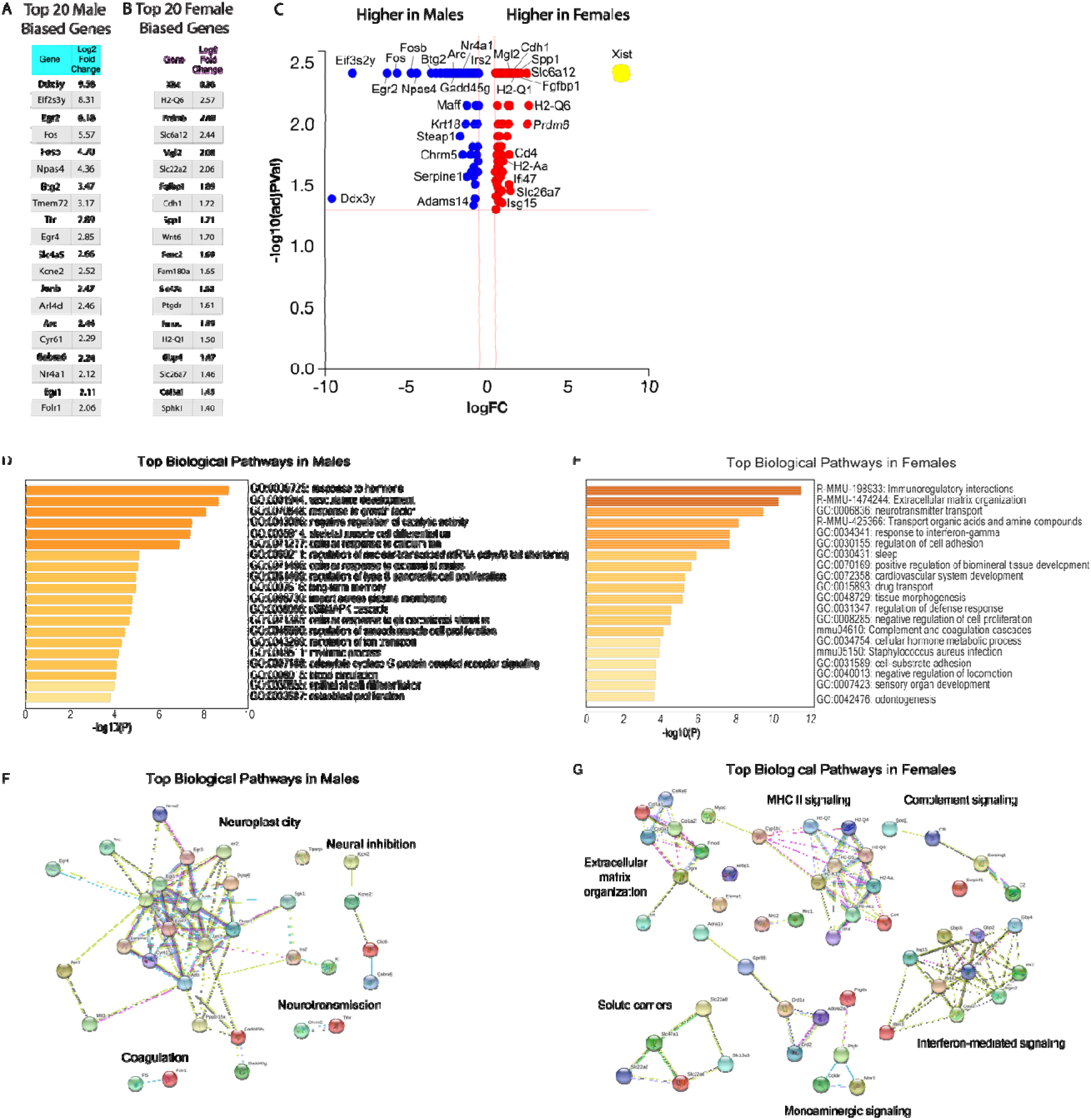
Differential gene expression in hippocampus of males *vs* females prior to immune challenge. (A,B) Top 10 differentially expressed (DEGs) in immune-naïve male *versus* female mice. 95 genes were more highly expressed in male hippocampi whereas 125 genes were more highly expressed in females. (C) Volcano plots showing genes that are higher in males (blue) and higher in females (red) at baseline obtained from Advaita iPathway Analysis. Differentially expressed (DE) genes are represented in terms of their measured expression change (x-axis) and the significance of the change (y-axis), with upregulated genes shown in red and downregulated genes shown in blue. (D,E) Biological pathways and processes enriched in (D) males compared with females and (E) in females compared with males. (F,G) Protein interaction (PPI) networks of targets, clustered by biological function (F) in males and (G) in females.

Biological processes for male-biased gene expression included response to hormone (log10(P) = −9.1), response to growth factor (log10(P) = −8.1), negative regulation of catalytic activity (log10(P) = −7.5), cellular response to calcium ion (log10(P) = −6.9), and negative regulation of nuclear transcribed mRNA poly(A) tail shortening (log10(P) = −5.1) (Figure 3D). Female-biased gene expression included biological processes such as extracellular matrix organization (log10(P) = −10.1), neurotransmitter transport (log10(P) = −9.3), response to interferon gamma (log10(P) = −7.5), cell adhesion (log10(P) = −7.5), drug transport (log10(P) = −5.2), regulation of defense response (log10(P) = −4.5), complement and coagulation cascades (log10(P) = −4.1), and cellular metabolic process (log10(P) = −3.9) (Figure 3E).

Protein-protein interaction analysis (STRING) revealed that targets more strongly expressed in the hippocampus of males are involved in G protein and calcium signaling, protein phosphorylation, as well as immediate early genes and DNA-methylation modifiers (Figure 3F). Targets that more strongly expressed in the hippocampus of females included extracellular matrix receptor genes, growth factor, neurotransmitter receptors, neurotransmitter transporter, protein phosphorylation, immune signaling, and complement-associated genes (Figure 3G).

To determine the contribution of initial sex differences in gene expression on the sex-specific changes after subchronic immune challenge, we examine the impact of prior immune challenge on genes more strongly expressed in males at baseline (“male-biased” genes) and those more strongly expressed in females (“female-biased” genes). Overall, in males, male-biased genes tended to be downregulated and female-biased genes upregulated months after immune challenge. Here, of the 46 genes higher in males at baseline, 34 were downregulated and only 12 upregulated; and of the 42 genes higher in females at baseline 41 were upregulated in males and only one downregulated long after subchronic immune challenge. Conversely, female-biased genes tended to be downregulated (13 of 13) and male-biased genes were more likely to be upregulated (3 of 3) in females after subchronic immune challenge. Specifically, the male-biased genes downregulated in males long after immune challenge included plasticity-related genes (*Arc,Npas4, Dusp1*). Female-biased genes upregulated in males included those associated with solute carriers (e.g. *Slc6a12, Slc22a6, Sphk1*) and extracellular matrix organization (e.g. *Bmp7, Col9a2*) (Supplementary Table 7).

In the Long-term+Acute group, female-biased genes were more likely to be upregulated in males and downregulated in females. Interestingly, whereas female-biased genes were strongly differentially expressed in both sexes, male-biased genes were not differentially expressed in either sex. In males, 7 of 7 female-biased genes were upregulated. In females, 53 of 75 were downregulated, including targets related to MHC class II (e.g. *H2-Aa, Cd74, Spp1*), cellular hormone metabolism (e.g. *Bmp6, Ttr*), and extracellular matrix organization (e.g. *Cdh1, Col8a1, Enpp2*) (Supplementary Table 8).

### 3.5 Shared targets and pathways amongst experimental conditions

We conducted meta-analyses for both males and females to examine similarities and differences between long-term and long-term + acute transcriptional changes in the hippocampus. In males, only one DEG was common to both Long-term condition and animals in the Long-term+Acute, demonstrating that the DEGs that are altered at baseline, and those that are differentially regulated in response to another immune challenge reflect different pathways or processes (Figure 4A). In females, only 1/26 DEGs in the Long-term condition were also dysregulated after dysregulated in the Long-term+Acute conditions suggesting that baseline changes in gene expression likely contribute to dysregulation of subsequent neuroimmune response (Figure 4C).

**Figure 4.**
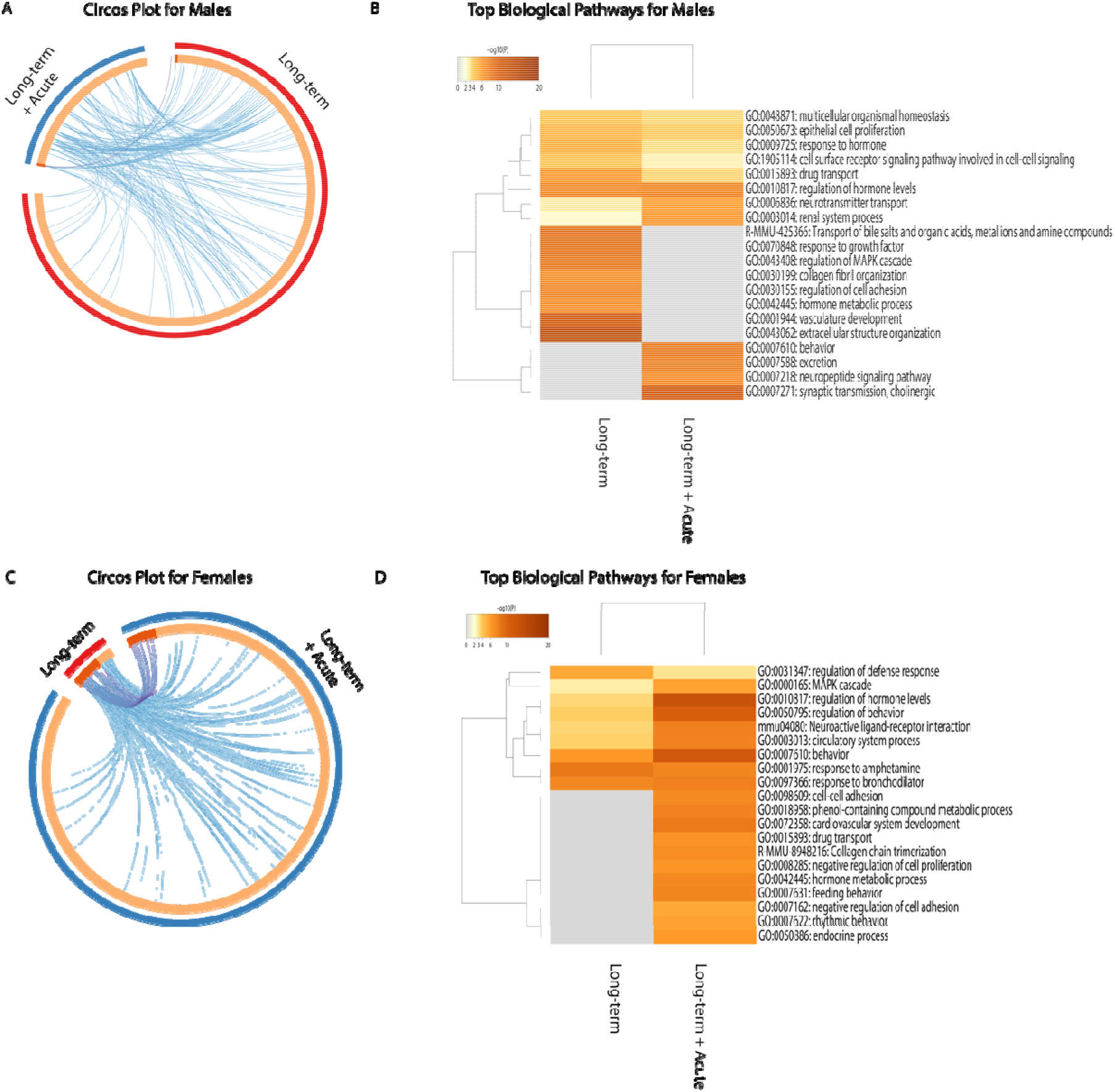
Meta-analysis of differentially expressed genes amongst long-term and acute conditions in males and females. (A) Males. Circos plot of differentially expressed genes amongst Long-term (3 months after subchronic LPS injections) and Long-term+Acute (LPS injection 3 months after subchronic challenge) conditions. DEG links shared between experimental conditions are depicted by purple lines while different genes that share similar biological pathways are depicted in blue. (B) Males. Heatmap of selected enriched gene ontology (GO) terms compared between Long-term and Long-term + Acute conditions in males. (C) Females. Circos plot of differentially expressed genes amongst Long-term and Long-term+Acute conditions. (D) Heatmap of selected enriched gene ontology (GO) terms compared between Long-term and Long-term + Acute conditions in females.

This analysis also compares biological pathways and processes between groups (blue lines). In males, only 3 biological pathways are shared, again demonstrating that baseline changes in gene expression long after immune challenge regulate processes other than response to acute challenge (Figure 4B). Unlike males, females showed a strong overlap between Long-Term and Long-term+Acute conditions, with 9 overlapping pathways, demonstrating the strongly increased transcriptional response to an acute challenge after prior immune experience. (Figure 4D).

Comparing these males and females in these analyses demonstrates the diverging impact of prior immune challenge on different components of immune function. Whereas males show persistent alterations in baseline expression of immune-related genes also activated during acute immune challenge, females show a fewer DEGs and less overlap with acute immune regulation. In contrast, males show little impact of a prior immune challenge on the acute transcriptional response in the hippocampus, whereas females that have experienced a prior immune challenge show a grossly exaggerated response to an acute inflammation.

## 4. DISCUSSION

Here we demonstrated long-lasting consequences of subchronic immune challenge, and striking sex differences, on gene expression in the hippocampus. In males, we observed enduring dysregulation of gene expression months after the end of a two-week immune challenge. In contrast, females showed few persistent changes at baseline, but striking changes in gene expression in response to an additional acute LPS injection three months after the subchronic immune challenge.

Males and females differed in both the magnitude of gene expression changes, and in the specific genes and pathways differentially expressed in the hippocampus long after immune challenge. In males but not females, we observed clear changes in both immune-related pathways and in genes and clusters related to neural plasticity. Immune-related pathways and genes, including response to interferon-gamma, complement signaling, and MHC II signaling were significantly dysregulated in males, and neuroplasticity-related clusters included extracellular matrix organization and cell adhesion molecules that are important for synaptic organization, as well as immediate early genes that are involved in activity-dependent transcriptional changes necessary for synaptic plasticity (Hawk and Abel, 2011; Minatohara et al., 2016). Several genes (e.g., *Fos, NR4a1/2, Egr, Spp1, Dusp1/6,Arc*) and pathways (MAPK signaling) are notably required for both immune- and neuroplasticity-related functions (Rosi, 2011; Donzis and Tronson, 2014; Stephen et al., 2017). In contrast, the dominant changes observed in females were pathways and genes related to monoaminergic signaling and its regulation, including adenosine, dopamine, and adrenergic receptors. Given the importance of dopaminergic and adrenergic signaling for memory modulation, motivational processes, and affective responses (Stone et al., 1999; Strange and Dolan, 2004; Badgaiyan et al., 2010; Wassum et al., 2011).

We also observed sex-specific transcriptional changes in the hippocampus in response to a subsequent immune challenge. Here, females previously exposed to a subchronic immune challenge showed greater dysregulation of transcriptional response after an acute LPS injection, whereas males showed similar patterns of gene expression whether or not they had previously experienced a subchronic challenge. Thus both males and females show changes in transcriptional processes in the brain that persist long after a subchronic immune challenge, but the patterns of dysregulation show striking sex specificity, and suggest that males and females have different patterns of vulnerability and resilience to future stressors, including immune challenge, and other environmental events (Tchessalova et al., 2018).

These sex-specific patterns of enduring changes in gene expression after subchronic immune challenge are consistent with strikingly different patterns of gene expression in the brain of males and females after stress or drugs of abuse (Hodes et al., 2015; Mychasiuk et al., 2016; Finn et al., 2018; Randesi et al., 2018), and further demonstrate that environmental insults have sex-specific implications for enduring changes in gene expression, neural function, and vulnerability to subsequent stressors. Such sex differences in transcriptional regulation may also provide new insights into differential vulnerability to cognitive decline, memory impairments, and affective disorders (Hogue et al., 2003; Liossi and Wood, 2014; Rainville and Hodes, 2019).

Sex-specific regulation of differential gene expression is consistent with sex differences in memory deficits observed in males and females after immune challenge or illness. We recently demonstrated that males but not females show deficits of fear memory, but that both sexes exhibit impairments of object recognition memory several months after subchronic immune challenge (Tchessalova and Tronson, 2019). Sex-specific patterns of memory deficits are also observed in patients, where women are more vulnerable to disruption of visuospatial tasks months after surgery or injury (Hogue et al., 2003; Liossi and Wood, 2014) whereas men show more progressive memory decline over the following years (Himanen et al., 2006). Determining the long-lasting changes in gene expression in males and in females is therefore important for identifying sex differences in the contribution of environmental insults to the vulnerability or resilience to cognitive decline, memory impairments, and affective disorders.

How baseline sex differences in hormonal levels (Koss and Frick, 2017; McEwen and Milner, 2017), memory processes (Keiser and Tronson, 2016; Keiser et al., 2017), emotion (Pitychoutis and Papadopoulou-Daifoti, 2010), and gene expression (Vied et al., 2016) mediate differential vulnerability to immune-triggered memory decline remains unknown. Here, we examined the relationship between sex-biased gene expression at baseline and differential regulation of those genes after immune challenge in males and females. We observed that most of the genes that were higher in the hippocampus of males at baseline were downregulated in the male hippocampus and upregulated in the female hippocampus long after immune challenge. Similarly, we found that female-biased genes were downregulated in females and upregulated in males after immune challenge. This pattern suggests that baseline differences in hippocampal gene expression contribute to their regulation after immune challenge. The pattern of regulation may be important for predicting which genes and pathways may be differentially vulnerable between the sexes to changes as a consequence of illness or stress.

There are a few limitations of these gene expression studies. First, in order to maintain RNA integrity, animals were not perfused prior to hippocampal dissections, and as such we cannot rule out persistent changes in gene expression from blood cells contributing to the changes in “hippocampal” gene expression observed here. Second, based on our previous work that demonstrated persistent memory deficits in both males and females after subchronic immune challenge, albeit fear conditioning deficits only males, we anticipated that males and females would show largely overlapping changings in gene expression, and our study was powered accordingly. As a result of the striking sex differences in DEG, we needed to consider males and females separately, resulting in relatively small sample size. Nevertheless, these findings provide important – and surprising – new data identifying persistent and sex-specific changes in transcriptional networks after subcrhonic immune challenge. Subsequent work will add to these findings using cell-type specific techniques across multiple brain regions to identify the causal roles of enduring changes in transcriptional regulation in long-lasting memory deficits after immune challenge.

These findings provide critical new insight into the long-lasting impact of immune-related signaling on gene expression in the brain. Understanding the long-lasting contributions of neuroimmune signaling to neuromodulation and plasticity is particularly important given the growing recognition that many environmental events, including stress (Weber et al., 2015; Wohleb et al., 2015; McKim et al., 2016; Frank et al., 2017; Serrats et al., 2017) and drugs of abuse (Crews et al., 2015; Hofford et al., 2018) both recruit neuroimmune signaling pathways. Here we demonstrated sex-specific patterns of gene expression months after a subchronic immune challenge, where males showed a persistent shift in baseline gene expression and females showed a markedly different response to subsequent stimulation. Enduring changes in genes and pathways that mediate plasticity-related processes, in addition to immune-related genes, in the hippocampus suggests that transient inflammatory signaling in the brain has important implications for neural function and hippocampal-dependent processes. As such, identifying the sex-specific changes in transcriptional regulation will be of importance for predicting vulnerabilities to and identifying novel, sex-specific biomarkers and therapeutic targets for cognitive decline, memory impairments, and affective disorders.

## Supporting information

Supplementary Table 1. Long-term DEGs in Males.

Supplementary Table 2. Long-term DEGs in Females.

Supplementary Table 3. Long-term + Acute DEGs in Males.

Supplementary Table 4. Long-term + Acute DEGs in Females.

Supplementary Table 5. Long-term + Acute shared between Males.and Females.

Supplementary Table 6. Male and Female Biased Genes.

Supplementary Table 7. Long-term_Male and Female Biased Genes.

Supplementary Table 8. Long-term + Acute_Male and Female Biased Genes.

## Acknowledgements

Thanks to University of Michigan Bioinformatics core for their expertise and data analysis of differentially expressed genes and the University of Michigan DNA sequencing core for their processing of samples; to Dr. Shigeki Iwase and Dr. Christina Vallianatos for assistance with prepping of the RNA samples; and Ms. Brynne Raines, Ms. Melanie Gil, and Ms. Caitlin Posillico for their extensive feedback on this manuscript

## REFERENCES

Adamsky A, Kol A, Kreisel T, Doron A, Ozeri-Engelhard N, Melcer T, Refaeli R, Horn H, Regev L, Groysman M, London M, Goshen I (2018) Astrocytic Activation Generates De Novo Neuronal Potentiation and Memory Enhancement. Cell 174:59–71.e14

Anderson ST, Commins S, Moynagh PN, Coogan AN (2015) Lipopolysaccharide-induced sepsis induces long-lasting affective changes in the mouse. Brain Behav Immun 43:98–109 Available at: http://linkinghub.elsevier.com/retrieve/pii/S0889159114003997

Avital A, Goshen I, Kamsler A, Segal M, Iverfeldt K, Richter-Levin G, Yirmiya R (2003) Impaired interleukin-1 signaling is associated with deficits in hippocampal memory processes and neural plasticity. Hippocampus 13:826–834

Badgaiyan RD, Fischman AJ, Alpert NM (2010) Dopamine Release During Human Emotional Processing. Neuroimage 47:2041–2045.

Bagot RC et al. (2016) Circuit-wide transcriptional profiling reveals brain region-specific gene networks regulating depression susceptibility Rosemary. Neuron 90:969–983.

Bekhbat M, Neigh GN (2017) Sex differences in the neuro-immune consequences of stress: Focus on depression and anxiety. Brain Behav Immun 67:1–12

Cates HM, Bagot RC, Heller EA, Purushothaman I, Lardner CK, Walker DM, Peña CJ, Neve RL, Shen L, Nestler EJ (2019) A novel role for E2F3b in regulating cocaine action in the prefrontal cortex. Neuropsychopharmacology 44:776–784.

Chavan SS, Huerta PT, Robbiati S, Valdes-Ferrer SI, Ochani M, Dancho M, Frankfurt M, Volpe BT, Tracey KJ, Diamond B (2012) HMGB1 mediates cognitive impairment in sepsis survivors. Mol Med 18:930–937.

Cholewa-Waclaw J, Bird A, von Schimmelmann M, Schaefer A, Yu H, Song H, Madabhushi R, Tsai L-H (2016) The Role of Epigenetic Mechanisms in the Regulation of Gene Expression in the Nervous System. J Neurosci 36:11427–11434

Cloutier CJ, Kavaliers M, Ossenkopp K-PP (2012) Lipopolysaccharide inhibits the simultaneous establishment of LiCl-induced anticipatory nausea and intravascularly conditioned taste avoidance in the rat. Behav Brain Res 232:278–286

Crews FT, John PD, Hill C, Carolina N, Sarkar DK, Ph D, Phil D (2015) Neuroimmune Function and the Consequences of Alcohol Exposure. Alcohol Res 37:331–351.

Dantzer R, Bluthé RM, Layé S, Bret-Dibat JL, Parnet P, Kelley KW (1998) Cytokines and sickness behavior. Ann N Y Acad Sci 840:586–590

del Rey A, Balschun D, Wetzel W, Randolf A, Besedovsky HO (2013) A cytokine network involving brain-borne IL-1β, IL-1ra, IL-18, IL-6, and TNFα operates during long-term potentiation and learning. Brain Behav Immun 33:15–23

Donzis EJ, Tronson NC (2014) Modulation of learning and memory by cytokines: signaling mechanisms and long term consequences. Neurobiol Learn Mem 115:68–77

Fenn AM, Gensel JC, Huang Y, Popovich PG, Lifshitz J, Godbout JP (2014) Immune Activation Promotes Depression 1 Month After Diffuse Brain Injury: A Role for Primed Microglia. Biol Psychiatry 76:575–584

Finn DA, Hashimoto JG, Cozzoli DK, Helms ML, Nipper MA, Kaufman MN, Wiren KM, Guizzetti M (2018) Binge Ethanol Drinking Produces Sexually Divergent and Distinct Changes in Nucleus Accumbens Signaling Cascades and Pathways in Adult C57BL/6J Mice. Front Genet 9:1–18.

Frank MG, Watkins LR, Maier SF (2017) Stress-induced glucocorticoids as a neuroendocrine alarm signal of danger. Brain Behav Immun 33:1–6.

Gharacholou SM, Reid KJ, Arnold S V, Rich MW, Pellikka PA, Singh M, Holsinger T, Krumholz HM, Peterson ED, Alexander KP, Clinic M (2011) Cognitive Impairment and Outcomes in Older Adult Survivors of Acute Myocardial Infarction: Findings from the TRIUMPH Registry. Am Hear J 162:860–869.

Gray J, Rubin T, Hunter R, McEwen B (2014) Hippocampal gene expression changes underlying stress sensitization and recovery. Mol Psychiatry 19:1171–1178.

Hawk JD, Abel T (2011) The role of NR4A transcription factors in memory formation. Brain Res Bull 85:21–29.

Himanen L, Portin R, Isoniemi H, Helenius H, Kurki T, Tenovuo O (2006) Longitudinal cognitive changes in traumatifc brain injury. Neurology 66:187–192.

Hodes GE et al. (2015) Sex Differences in Nucleus Accumbens Transcriptome Profiles Associated with Susceptibility versus Resilience to Subchronic Variable Stress. J Neurosci 35:16362–16376

Hofford RS, Russo SJ, Kiraly DD (2018) Neuroimmune mechanisms of psychostimulant and opioid use disorders. Eur J Neurosci:1–12.

Hogue CW, Lillie R, Hershey T, Birge S, Nassief AM, Thomas B, Freedland KE (2003) Gender Influence on Cognitive Function After Cardiac Operation. Annu Thorac Surg 4975.

Huerta PT, Robbiati S, Huerta TS, Sabharwal A, Berlin RA, Frankfurt M, Volpe BT (2016) Preclinical models of overwhelming sepsis implicate the neural system that encodes contextual fear memory. Mol Med 22.

Keiser AA, Tronson NC (2016) Molecular mechanisms of memory in males and females. In: Sex Differences in the Central Nervous System, 1st ed. (Shansky RM, ed), pp 27–46. Elsevier Inc.

Keiser AA, Turnbull LM, Darian MA, Feldman DE, Song I, Tronson NC (2017) Sex differences in context fear generalization and recruitment of hippocampus and amygdala during retrieval. Neuropsychopharmacology 42:397–407.

Kondo S, Kohsaka S, Okabe S (2011) Long-term changes of spine dynamics and microglia after transient peripheral immune response triggered by LPS in vivo. Mol Brain 4:27

Koss WA, Frick KM (2017) Sex differences in hippocampal function. J Neurosci Res 95:539–562

Liossi C, Wood RL (2014) Gender as a Moderator of Cognitive and Affective Outcome After Traumatic Brain Injury. J Neuropsychiatry Clin Neurosci 21:43–51.

Liu Y, Qin L, Wilson B, Wu X, Qian L, Granholm A-CC, Crews FT, Hong J-SS (2008) Endotoxin induces a delayed loss of TH-IR neurons in substantia nigra and motor behavioral deficits. Neurotoxicology 29:864–870.

Lorsch ZS et al. (2018) Estrogen receptor *α* drives pro-resilient transcription in mouse models of depression Zachary. Nat Commun 9:1–11.

Marra A, Pandharipande PP, Girard TD, Patel MB, Hughes CG, Jackson JC, Thompson JL, Chandrasekhar R, Ely EW, Brummel NE (2018) Co-Occurrence of Post-Intensive Care Syndrome Problems Among 406 Survivors of Critical Illness*. Crit Care Med 46:1393–1401.

McEwen BS, Milner TA (2017) Understanding the broad influence of sex hormones and sex differences in the brain. J Neurosci Res 95:24–39.

McKim DB, Niraula A, Tarr AJ, Wohleb ES, Sheridan JF, Godbout JP (2016) Neuroinflammatory Dynamics Underlie Memory Impairments after Repeated Social Defeat. J Neurosci 36:2590–2604.

Minatohara K, Akiyoshi M, Okuno H (2016) Role of Immediate-Early Genes in Synaptic Plasticity and Neuronal Ensembles Underlying the Memory Trace. Front Mol Neurosci 8.

Ming Z, Sawicki G, Bekar LK (2015) Acute systemic LPS-mediated inflammation induces lasting changes in mouse cortical neuromodulation and behavior. Neurosci Lett 590:96–100

Mychasiuk R, Muhammad A, Kolb B (2016) Chronic stress induces persistent changes in global DNA methylation and gene expression in the medial prefrontal cortex, orbitofrontal cortex, and hippocampus. Neuroscience 322:489–499.

Nestler EJ (2014) Epigenetic Mechanisms of Drug Addiction. Neuropharmacology 76:1–22.

Netea MG, van der Meer JWM (2017) Trained Immunity: An Ancient Way of Remembering. Cell Host Microbe 21:297–300

Norden DM, Muccigrosso MM, Godbout JP (2015) Microglial priming and enhanced reactivity to secondary insult in aging, and traumatic CNS injury, and neurodegenerative disease. Neuropharmacology 96:29–41

Pitychoutis PM, Papadopoulou-Daifoti Z (2010) Of depression and immunity: does sex matter? Int J Neuropsychopharmacol 13:675–689.

Rainville JR, Hodes GE (2019) Inflaming sex differences in mood disorders. Neuropsychopharmacology 44:184–199

Raison C, Capuron L, Miller A (2006) Cytokines sing the blues: inflammation and the pathogenesis of depression. Trends Immunol 27:24–31.

Randesi M, Zhou Y, Mazid S, Odell SC, Gray JD, Correa da Rosa J, McEwen BS, Milner TA, Kreek MJ (2018) Sex differences after chronic stress in the expression of opioid-and neuroplasticity-related genes in the rat hippocampus. Neurobiol Stress 8:33–41.

Rosi S (2011) Neuroinflammation and the plasticity-related immediate-early gene Arc. Brain Behav Immun 25:S39–S49

Semmler A, Frisch C, Debeir T, Ramanathan M, Okulla T, Klockgether T, Heneka MT (2007) Long-term cognitive impairment, neuronal loss and reduced cortical cholinergic innervation after recovery from sepsis in a rodent model. Exp Neurol 204:733–740.

Serrats J, Grigoleit JS, Alvarez-Salas E, Sawchenko PE (2017) Pro-inflammatory immune-to-brain signaling is involved in neuroendocrine responses to acute emotional stress. Brain Behav Immun 62:53–63.

Singer M, Deutschman CS, Seymour C, Shankar-Hari M, Annane D, Bauer M, Bellomo R, Bernard GR, Chiche JD, Coopersmith CM, Hotchkiss RS, Levy MM, Marshall JC, Martin GS, Opal SM, Rubenfeld GD, Poll T Der, Vincent JL, Angus DC (2016) The third international consensus definitions for sepsis and septic shock (sepsis-3). JAMA - J Am Med Assoc 315.

Stephen S, Jin U, Morpurgo B, Abdayyeh A, Singh M, Tjalkens RB (2017) Nuclear Receptor 4A (NR4A) Family – Orphans No More. J Steroid Biochem Mol Biol 4:48–60.

Sterlemann V, Ganea K, Liebl C, Harbich D, Alam S, Holsboer F, Müller MB, Schmidt M V. (2008) Long-term behavioral and neuroendocrine alterations following chronic social stress in mice: Implications for stress-related disorders. Horm Behav 53:386–394.

Stone EA, Zhang Y, Rosengarten H, Yeretsian J, Quartermain D (1999) Brain alpha 1-adrenergic neurotransmission is necessary for behavioral activation to environmental change in mice. Neuroscience 94:1245–1252.

Strange BA, Dolan RJ (2004) Adrenergic modulation of emotional memory-evoked human amygdala and hippocampal responses. Proc Natl Acad Sci 101:11454–11458.

Tafet GE, Nemeroff CB (2016) The Links Between Stress and Depression: Psychoneuroendocrinological, Genetic, and Environmental Interactions. J Neuropsychiatry Clin Neurosci 28:77–88.

Tchessalova D, Posillico CK, Tronson NC (2018) Neuroimmune activation drives multiple brain states. Front Syst Neurosci 12:39

Tchessalova D, Tronson NC (2019) Memory deficits in males and females long after subchronic immune challenge. Neurobiol Learn Mem 158:60–72

Vied C, Ray S, Badger C, Bundy JL, Arbeitman MN, Nowakowski RS (2016) Transcriptomic Analysis of the Hippocampus From Six Inbred Strains of Mice Suggests a Basis for Sex-Specific Susceptibility and Severity of Neurological Disorders. J Comp Neurol 524:2696–2710.

Volpe BT, Berlin RA, Frankfurt M (2015) The brain at risk: the sepsis syndrome and lessons from preclinical experiments. Immunol Res 63:70–74

Walker DM et al. (2018) Cocaine Self-administration Alters Transcriptome-wide Responses in the Brain’s Reward Circuitry. Biol Psychiatry 84:867–880.

Wassum KM, Ostlund SB, Balleine BW, Maidment NT (2011) Differential dependence of Pavlovian incentive motivation and instrumental incentive learning processes on dopamine signaling. Learn Mem 18:475–483.

Weber MD, Frank MG, Tracey KJ, Watkins LR, Maier SF (2015) Stress Induces the Danger-Associated Molecular Pattern HMGB-1 in the Hippocampus of Male Sprague Dawley Rats: A Priming Stimulus of Microglia and the NLRP3 Inflammasome. J Neurosci 35:316–324.

Weberpals M, Hermes M, Hermann S, Kummer MP, Terwel D, Semmler A, Berger M, Schafers M, Heneka MT, Schäfers M, Heneka MT (2009) NOS2 gene deficiency protects from sepsis-induced long-term cognitive deficits. J Neurosci 29:14177–14184.

Wendeln A-C et al. (2018) Innate immune memory in the brain shapes neurological disease hallmarks. Nature 556:332–338

Wohleb ES, McKim DB, Sheridan JF, Godbout JP (2015) Monocyte trafficking to the brain with stress and inflammation: A novel axis of immune-to-brain communication that influences mood and behavior. Front Neurosci 9:1–17.

Yirmiya R, Goshen I (2011) Immune modulation of learning, memory, neural plasticity and neurogenesis. Brain Behav Immun 25:181–213

