## Supplementary Table 1. Long-term DEGs in Males. for "Enduring and sex-specific changes in hippocampal gene expression after a subchronic immune challenge"

**Supplementary Table 1. Differentially expressed genes in the hippocampus of males three months after subchronic immune challenge (Long-term condition females).**

| **Gene** | **Gene Name** | **Log2 Fold Change** | **P-value** |
| --- | --- | --- | --- |
| *Wfdc2* | WAP four-disulfide core domain 2 | 2.37 | 0.004 |
| *Ttr* | transthyretin | 2.30 | 0.004 |
| *Aqp1* | aquaporin 1 | 2.25 | 0.004 |
| *Steap1* | six transmembrane epithelial antigen of the prostate 1 | 2.23 | 0.004 |
| *Prdm6* | PR domain containing 6 | 2.22 | 0.022 |
| *Slc16a8* | solute carrier family 16 (monocarboxylic acid transporters), member 8 | 2.22 | 0.004 |
| *Tmem72* | transmembrane protein 72 | 2.22 | 0.004 |
| *Cldn2* | claudin 2 | 2.18 | 0.004 |
| *Wdr72* | WD repeat domain 72 | 2.17 | 0.004 |
| *Kcne2* | potassium voltage-gated channel, Isk-related subfamily, gene 2 | 2.16 | 0.004 |
| *1500015O10Rik* | RIKEN cDNA 1500015O10 gene | 2.14 | 0.004 |
| *Cldn9* | claudin 9 | 2.09 | 0.040 |
| *Clic6* | chloride intracellular channel 6 | 2.08 | 0.004 |
| *Slc6a12* | solute carrier family 6 (neurotransmitter transporter, betaine/GABA), member 12 | 2.06 | 0.004 |
| *Tmprss11a* | transmembrane protease, serine 11a | 2.05 | 0.004 |
| *1110059M19Rik* | proline rich 32 | 2.05 | 0.004 |
| *Slc4a5* | solute carrier family 4, sodium bicarbonate cotransporter, member 5 | 2.04 | 0.004 |
| *Sptlc3* | serine palmitoyltransferase, long chain base subunit 3 | 2.03 | 0.004 |
| *Folr1* | folate receptor 1 (adult) | 2.03 | 0.004 |
| *Oca2* | oculocutaneous albinism II | 2.00 | 0.004 |
| *Sult1c2* | sulfotransferase family, cytosolic, 1C, member 2 | 1.99 | 0.004 |
| *Wdr86* | WD repeat domain 86 | 1.98 | 0.004 |
| *Defb11* | defensin beta 11 | 1.91 | 0.040 |
| *Slco1a5* | solute carrier organic anion transporter family, member 1a5 | 1.90 | 0.004 |
| *F5* | coagulation factor V | 1.88 | 0.004 |
| *Tmem27* | transmembrane protein 27 | 1.88 | 0.024 |
| *Sostdc1* | sclerostin domain containing 1 | 1.87 | 0.004 |
| *Tc2n* | tandem C2 domains, nuclear | 1.85 | 0.007 |
| *Tmem184a* | transmembrane protein 184a | 1.83 | 0.007 |
| *Col8a1* | collagen, type VIII, alpha 1 | 1.82 | 0.004 |
| *Gyltl1b* | glycosyltransferase-like 1B | 1.80 | 0.004 |
| *Scara5* | scavenger receptor class A, member 5 | 1.78 | 0.004 |
| *Col17a1* | collagen, type XVII, alpha 1 | 1.77 | 0.004 |
| *Enpp2* | ectonucleotide pyrophosphatase/phosphodiesterase 2 | 1.76 | 0.004 |
| *Rrh* | retinal pigment epithelium derived rhodopsin homolog | 1.76 | 0.040 |
| *Pla2g5* | phospholipase A2, group V | 1.73 | 0.004 |
| *Otx2* | orthodenticle homeobox 2 | 1.73 | 0.004 |
| *Prlr* | prolactin receptor | 1.71 | 0.004 |
| *Rab20* | RAB20, member RAS oncogene family | 1.70 | 0.004 |
| *Krt18* | keratin 18 | 1.69 | 0.004 |
| *Cdh3* | cadherin 3 | 1.67 | 0.015 |
| *Trpv4* | transient receptor potential cation channel, subfamily V, member 4 | 1.65 | 0.004 |
| *Slc26a7* | solute carrier family 26, member 7 | 1.65 | 0.018 |
| *Sema3b* | sema domain, immunoglobulin domain (Ig), short basic domain, secreted, (semaphorin) 3B | 1.64 | 0.004 |
| *Foxc2* | forkhead box C2 | 1.62 | 0.004 |
| *Ace* | angiotensin I converting enzyme (peptidyl-dipeptidase A) 1 | 1.60 | 0.004 |
| *Kl* | klotho | 1.58 | 0.004 |
| *Slc13a4* | solute carrier family 13 (sodium/sulfate symporters), member 4 | 1.57 | 0.004 |
| *Abca4* | ATP-binding cassette, sub-family A (ABC1), member 4 | 1.57 | 0.004 |
| *Col8a2* | collagen, type VIII, alpha 2 | 1.56 | 0.004 |
| *Ccdc135* | dynein regulatory complex subunit 7 | 1.56 | 0.004 |
| *4833427G06Rik* | RIKEN cDNA 4833427G06 gene | 1.56 | 0.037 |
| *Krt8* | keratin 8 | 1.56 | 0.004 |
| *Sulf1* | sulfatase 1 | 1.56 | 0.004 |
| *Igf2* | insulin-like growth factor 2 | 1.55 | 0.004 |
| *Krt23* | keratin 23 | 1.54 | 0.035 |
| *Col4a3* | collagen, type IV, alpha 3 | 1.53 | 0.004 |
| *Wfikkn2* | WAP, follistatin/kazal, immunoglobulin, kunitz and netrin domain containing 2 | 1.49 | 0.004 |
| *Rdh5* | retinol dehydrogenase 5 | 1.49 | 0.004 |
| *Sfrp5* | secreted frizzled-related sequence protein 5 | 1.48 | 0.004 |
| *Lmx1a* | LIM homeobox transcription factor 1 alpha | 1.46 | 0.004 |
| *Slc22a2* | solute carrier family 22 (organic cation transporter), member 2 | 1.45 | 0.035 |
| *Plek2* | pleckstrin 2 | 1.45 | 0.040 |
| *Cdh1* | cadherin 1 | 1.44 | 0.004 |
| *Tcea3* | transcription elongation factor A (SII), 3 | 1.44 | 0.020 |
| *Pcolce* | procollagen C-endopeptidase enhancer protein | 1.39 | 0.004 |
| *Lbp* | lipopolysaccharide binding protein | 1.37 | 0.004 |
| *Wnt6* | wingless-type MMTV integration site family, member 6 | 1.37 | 0.010 |
| *Rbm47* | RNA binding motif protein 47 | 1.36 | 0.004 |
| *Pon3* | paraoxonase 3 | 1.34 | 0.004 |
| *Tuba1c* | tubulin, alpha 1C | 1.33 | 0.018 |
| *Igfbp2* | insulin-like growth factor binding protein 2 | 1.33 | 0.004 |
| *A2m* | alpha-2-macroglobulin | 1.32 | 0.004 |
| *Six3os1* | SIX homeobox 3, opposite strand 1 | 1.29 | 0.048 |
| *Fgfbp1* | fibroblast growth factor binding protein 1 | 1.29 | 0.022 |
| *Epn3* | epsin 3 | 1.29 | 0.004 |
| *Cndp1* | carnosine dipeptidase 1 (metallopeptidase M20 family) | 1.26 | 0.026 |
| *Fam180a* | family with sequence similarity 180, member A | 1.24 | 0.007 |
| *Col4a4* | collagen, type IV, alpha 4 | 1.24 | 0.004 |
| *Col9a3* | collagen, type IX, alpha 3 | 1.23 | 0.004 |
| *H2-Q1* | histocompatibility 2, Q region locus 1 | 1.22 | 0.037 |
| *Ptgds* | prostaglandin D2 synthase (brain) | 1.22 | 0.004 |
| *Slc39a4* | solute carrier family 39 (zinc transporter), member 4 | 1.20 | 0.004 |
| *Cdkn1c* | cyclin-dependent kinase inhibitor 1C (P57) | 1.20 | 0.004 |
| *Slc47a1* | solute carrier family 47, member 1 | 1.19 | 0.010 |
| *H2-Ab1* | histocompatibility 2, class II antigen A, beta 1 | 1.17 | 0.004 |
| *Spp1* | secreted phosphoprotein 1 | 1.16 | 0.004 |
| *Fmod* | fibromodulin | 1.15 | 0.004 |
| *Arhgap28* | Rho GTPase activating protein 28 | 1.15 | 0.031 |
| *Sphk1* | sphingosine kinase 1 | 1.15 | 0.004 |
| *Ptgdr* | prostaglandin D receptor | 1.14 | 0.026 |
| *Fap* | fibroblast activation protein | 1.14 | 0.007 |
| *Gpr81* | hydrocarboxylic acid receptor 1 | 1.13 | 0.018 |
| *Trpm3* | transient receptor potential cation channel, subfamily M, member 3 | 1.13 | 0.004 |
| *H2-Aa* | histocompatibility 2, class II antigen A, alpha | 1.11 | 0.004 |
| *Serpinb1b* | serine (or cysteine) peptidase inhibitor, clade B, member 1b | 1.11 | 0.013 |
| *Gpx8* | glutathione peroxidase 8 (putative) | 1.10 | 0.004 |
| *H2-Eb1* | histocompatibility 2, class II antigen E beta | 1.10 | 0.013 |
| *Tgfbi* | transforming growth factor, beta induced | 1.10 | 0.004 |
| *Crabp2* | cellular retinoic acid binding protein II | 1.09 | 0.004 |
| *Cd59a* | CD59a antigen | 1.09 | 0.004 |
| *Mdfic* | MyoD family inhibitor domain containing | 1.09 | 0.004 |
| *Msx1as* | msh homeobox 1 opposite strand | 1.09 | 0.004 |
| *Msx1* | msh homeobox 1 | 1.08 | 0.029 |
| *Aldh1a2* | aldehyde dehydrogenase family 1, subfamily A2 | 1.08 | 0.004 |
| *Cd74* | CD74 antigen (invariant polypeptide of major histocompatibility complex, class II antigen-associated) | 1.08 | 0.004 |
| *Glb1l2* | galactosidase, beta 1-like 2 | 1.07 | 0.004 |
| *Vat1l* | vesicle amine transport protein 1 like | 1.07 | 0.004 |
| *Cldn1* | claudin 1 | 1.06 | 0.004 |
| *Sdc1* | syndecan 1 | 1.04 | 0.048 |
| *Col3a1* | collagen, type III, alpha 1 | 1.04 | 0.004 |
| *Slc37a2* | solute carrier family 37 (glycerol-3-phosphate transporter), member 2 | 1.04 | 0.004 |
| *Bmp6* | bone morphogenetic protein 6 | 1.04 | 0.004 |
| *Slc22a6* | solute carrier family 22 (organic anion transporter), member 6 | 1.03 | 0.004 |
| *Steap2* | six transmembrane epithelial antigen of prostate 2 | 1.03 | 0.004 |
| *Slc6a13* | solute carrier family 6 (neurotransmitter transporter, GABA), member 13 | 1.02 | 0.004 |
| *Itih2* | inter-alpha trypsin inhibitor, heavy chain 2 | 1.01 | 0.004 |
| *Slc6a20a* | solute carrier family 6 (neurotransmitter transporter), member 20A | 1.00 | 0.004 |
| *Perp* | PERP, TP53 apoptosis effector | 1.00 | 0.004 |
| *Mpzl2* | myelin protein zero-like 2 | 0.99 | 0.010 |
| *Col6a4* | collagen, type VI, alpha 4 | 0.99 | 0.004 |
| *C2* | complement component 2 (within H-2S) | 0.98 | 0.010 |
| *Col1a1* | collagen, type I, alpha 1 | 0.98 | 0.004 |
| *Bmp7* | bone morphogenetic protein 7 | 0.97 | 0.004 |
| *Islr* | immunoglobulin superfamily containing leucine-rich repeat | 0.97 | 0.004 |
| *Slc16a9* | solute carrier family 16 (monocarboxylic acid transporters), member 9 | 0.96 | 0.004 |
| *Ppp1r3b* | protein phosphatase 1, regulatory (inhibitor) subunit 3B | 0.94 | 0.007 |
| *C1qtnf5* | C1q and tumor necrosis factor related protein 5 | 0.93 | 0.015 |
| *Itpripl1* | inositol 1,4,5-triphosphate receptor interacting protein-like 1 | 0.93 | 0.007 |
| *Serping1* | serine (or cysteine) peptidase inhibitor, clade G, member 1 | 0.92 | 0.004 |
| *Bst2* | bone marrow stromal cell antigen 2 | 0.91 | 0.004 |
| *Col4a6* | collagen, type IV, alpha 6 | 0.90 | 0.033 |
| *Slc4a2* | solute carrier family 4 (anion exchanger), member 2 | 0.90 | 0.004 |
| *Olfml2a* | olfactomedin-like 2A | 0.90 | 0.026 |
| *Ogn* | osteoglycin | 0.88 | 0.010 |
| *Aox3* | aldehyde oxidase 3 | 0.88 | 0.020 |
| *Col1a2* | collagen, type I, alpha 2 | 0.87 | 0.004 |
| *Crhr2* | corticotropin releasing hormone receptor 2 | 0.85 | 0.031 |
| *Lgals3bp* | lectin, galactoside-binding, soluble, 3 binding protein | 0.85 | 0.004 |
| *Ifit1* | interferon-induced protein with tetratricopeptide repeats 1 | 0.84 | 0.029 |
| *Cgnl1* | cingulin-like 1 | 0.83 | 0.004 |
| *Car12* | carbonic anhydrase 12 | 0.83 | 0.004 |
| *Gstm2* | glutathione S-transferase, mu 2 | 0.83 | 0.039 |
| *Emilin1* | elastin microfibril interfacer 1 | 0.82 | 0.037 |
| *Mrc1* | mannose receptor, C type 1 | 0.81 | 0.010 |
| *Ltc4s* | leukotriene C4 synthase | 0.80 | 0.039 |
| *Oasl2* | 2'-5' oligoadenylate synthetase-like 2 | 0.80 | 0.031 |
| *Rbp1* | retinol binding protein 1, cellular | 0.79 | 0.004 |
| *Slc16a12* | solute carrier family 16 (monocarboxylic acid transporters), member 12 | 0.79 | 0.004 |
| *Slc13a3* | solute carrier family 13 (sodium-dependent dicarboxylate transporter), member 3 | 0.78 | 0.004 |
| *Gprc5c* | G protein-coupled receptor, family C, group 5, member C | 0.78 | 0.018 |
| *Mrc2* | mannose receptor, C type 2 | 0.77 | 0.004 |
| *Sfrp1* | secreted frizzled-related protein 1 | 0.76 | 0.004 |
| *Aebp1* | AE binding protein 1 | 0.75 | 0.004 |
| *Cab39l* | calcium binding protein 39-like | 0.75 | 0.004 |
| *Csrp2* | cysteine and glycine-rich protein 2 | 0.73 | 0.040 |
| *Arhgef5* | Rho guanine nucleotide exchange factor (GEF) 5 | 0.72 | 0.044 |
| *Gjb2* | gap junction protein, beta 2 | 0.72 | 0.004 |
| *Mgp* | matrix Gla protein | 0.72 | 0.004 |
| *Slc31a1* | solute carrier family 31, member 1 | 0.72 | 0.004 |
| *Sgms2* | sphingomyelin synthase 2 | 0.72 | 0.013 |
| *Ifitm3* | interferon induced transmembrane protein 3 | 0.70 | 0.004 |
| *Vamp8* | vesicle-associated membrane protein 8 | 0.69 | 0.004 |
| *Coch* | cochlin | 0.69 | 0.007 |
| *Fbln1* | fibulin 1 | 0.69 | 0.004 |
| *Lum* | lumican | 0.69 | 0.018 |
| *Lepr* | leptin receptor | 0.69 | 0.029 |
| *Nqo1* | NAD(P)H dehydrogenase, quinone 1 | 0.68 | 0.022 |
| *Hfe* | hemochromatosis | 0.68 | 0.035 |
| *Pcolce2* | procollagen C-endopeptidase enhancer 2 | 0.68 | 0.010 |
| *Myof* | myoferlin | 0.68 | 0.013 |
| *Efemp1* | epidermal growth factor-containing fibulin-like extracellular matrix protein 1 | 0.66 | 0.004 |
| *Pgcp* | carboxypeptidase Q | 0.66 | 0.004 |
| *Dab2* | disabled 2, mitogen-responsive phosphoprotein | 0.65 | 0.004 |
| *Tcn2* | transcobalamin 2 | 0.64 | 0.004 |
| *Htr2c* | 5-hydroxytryptamine (serotonin) receptor 2C | 0.63 | 0.004 |
| *Mxra8* | matrix-remodelling associated 8 | 0.63 | 0.020 |
| *Lyz2* | lysozyme 2 | 0.61 | 0.024 |
| *Myoc* | myocilin | 0.60 | 0.004 |
| *Col9a2* | collagen, type IX, alpha 2 | 0.60 | 0.042 |
| *Lgals1* | lectin, galactose binding, soluble 1 | 0.60 | 0.024 |
| *Ucp2* | uncoupling protein 2 (mitochondrial, proton carrier) | 0.60 | 0.004 |
| *Sned1* | sushi, nidogen and EGF-like domains 1 | 0.60 | 0.042 |
| *Cfh* | complement component factor h | 0.60 | 0.004 |
| *Csrnp1* | cysteine-serine-rich nuclear protein 1 | -0.59 | 0.026 |
| *Arl5b* | ADP-ribosylation factor-like 5B | -0.63 | 0.007 |
| *Nfkbiz* | nuclear factor of kappa light polypeptide gene enhancer in B cells inhibitor, zeta | -0.64 | 0.004 |
| *Car8* | carbonic anhydrase 8 | -0.66 | 0.022 |
| *Per1* | period circadian clock 1 | -0.67 | 0.004 |
| *Gadd45b* | growth arrest and DNA-damage-inducible 45 beta | -0.75 | 0.004 |
| *Klf4* | Kruppel-like factor 4 (gut) | -0.78 | 0.004 |
| *Dusp6* | dual specificity phosphatase 6 | -0.78 | 0.004 |
| *Ppp1r15a* | protein phosphatase 1, regulatory (inhibitor) subunit 15A | -0.86 | 0.004 |
| *Nfil3* | nuclear factor, interleukin 3, regulated | -0.90 | 0.004 |
| *Adamts1* | a disintegrin-like and metallopeptidase (reprolysin type) with thrombospondin type 1 motif, 1 | -0.90 | 0.004 |
| *Bub1b* | BUB1B, mitotic checkpoint serine/threonine kinase | -0.91 | 0.004 |
| *Cenpa* | centromere protein A | -0.95 | 0.018 |
| *Tiparp* | TCDD-inducible poly(ADP-ribose) polymerase | -1.03 | 0.004 |
| *Rgs2* | regulator of G-protein signaling 2 | -1.04 | 0.004 |
| *Rasl11a* | RAS-like, family 11, member A | -1.09 | 0.004 |
| *Pcdh8* | protocadherin 8 | -1.12 | 0.004 |
| *Fosl2* | fos-like antigen 2 | -1.14 | 0.004 |
| *Sik1* | salt inducible kinase 1 | -1.20 | 0.004 |
| *Zfp36* | zinc finger protein 36 | -1.27 | 0.004 |
| *Apold1* | apolipoprotein L domain containing 1 | -1.28 | 0.004 |
| *Capn11* | calpain 11 | -1.35 | 0.004 |
| *Nr4a2* | nuclear receptor subfamily 4, group A, member 2 | -1.38 | 0.004 |
| *Cbln3* | cerebellin 3 precursor protein | -1.45 | 0.004 |
| *Gm13889* | predicted gene 13889 | -1.46 | 0.004 |
| *Errfi1* | ERBB receptor feedback inhibitor 1 | -1.48 | 0.004 |
| *Egr3* | early growth response 3 | -1.51 | 0.004 |
| *Kcnj2* | potassium inwardly-rectifying channel, subfamily J, member 2 | -1.56 | 0.004 |
| *Dusp1* | dual specificity phosphatase 1 | -1.61 | 0.004 |
| *Ptgs2* | prostaglandin-endoperoxide synthase 2 | -1.63 | 0.004 |
| *Gadd45g* | growth arrest and DNA-damage-inducible 45 gamma | -1.64 | 0.004 |
| *Ier2* | immediate early response 2 | -1.74 | 0.004 |
| *Atf3* | activating transcription factor 3 | -1.80 | 0.004 |
| *Egr1* | early growth response 1 | -1.91 | 0.004 |
| *Nr4a1* | nuclear receptor subfamily 4, group A, member 1 | -1.96 | 0.004 |
| *Arc* | activity regulated cytoskeletal-associated protein | -2.06 | 0.004 |
| *Ccl3* | chemokine (C-C motif) ligand 3 | -2.22 | 0.031 |
| *Junb* | jun B proto-oncogene | -2.28 | 0.004 |
| *Cyr61* | cysteine rich protein 61 | -2.30 | 0.004 |
| *Arl4d* | ADP-ribosylation factor-like 4D | -2.43 | 0.004 |
| *Egr4* | early growth response 4 | -2.60 | 0.004 |
| *Btg2* | B cell translocation gene 2, anti-proliferative | -3.20 | 0.004 |
| *Npas4* | neuronal PAS domain protein 4 | -3.96 | 0.004 |
| *Fos* | FBJ osteosarcoma oncogene | -4.76 | 0.004 |
| *Fosb* | FBJ osteosarcoma oncogene B | -4.81 | 0.004 |
| *Egr2* | early growth response 2 | -4.92 | 0.004 |
