## Supplementary Table 2. Long-term DEGs in Females. for "Enduring and sex-specific changes in hippocampal gene expression after a subchronic immune challenge"

**Supplementary Table 2. Differentially expressed genes in the hippocampus of females three months after subchronic immune challenge (Long-term condition females).**

| **Gene** | **Gene Name** | **Log2 Fold Change** | **P-value** |
| --- | --- | --- | --- |
| *Npas4* | neuronal PAS domain protein 4 | 1.27 | 0.004 |
| *Dio3* | deiodinase, iodothyronine type III | 1.14 | 0.007 |
| *Mid1* | midline 1 | 0.87 | 0.033 |
| *Fos* | FBJ osteosarcoma oncogene | 0.84 | 0.004 |
| *Gpr101* | G protein-coupled receptor 101 | 0.82 | 0.013 |
| *1700048O20Rik* | RIKEN cDNA 1700048O20 gene | 0.79 | 0.004 |
| *Dlk1* | delta-like 1 homolog (Drosophila) | 0.73 | 0.033 |
| *Cckbr* | cholecystokinin B receptor | -0.59 | 0.004 |
| *Coch* | cochlin | -0.61 | 0.013 |
| *Irgm2* | immunity-related GTPase family M member 2 | -0.64 | 0.048 |
| *Spp1* | secreted phosphoprotein 1 | -0.68 | 0.004 |
| *Ppp1r1b* | protein phosphatase 1, regulatory (inhibitor) subunit 1B | -0.69 | 0.004 |
| *Rasgef1c* | RasGEF domain family, member 1C | -0.70 | 0.020 |
| *Gbp5* | guanylate binding protein 5 | -0.70 | 0.046 |
| *Adra1b* | adrenergic receptor, alpha 1b | -0.76 | 0.013 |
| *Scn4b* | sodium channel, type IV, beta | -0.76 | 0.004 |
| *Myl4* | myosin, light polypeptide 4 | -0.82 | 0.042 |
| *Foxp2* | forkhead box P2 | -0.82 | 0.013 |
| *Ifit1* | interferon-induced protein with tetratricopeptide repeats 1 | -0.85 | 0.018 |
| *Gpr88* | G-protein coupled receptor 88 | -0.88 | 0.004 |
| *Gm7120* | predicted gene 7120 | -0.98 | 0.004 |
| *Gbp4* | guanylate binding protein 4 | -1.02 | 0.004 |
| *Drd2* | dopamine receptor D2 | -1.10 | 0.013 |
| *Cd4* | CD4 antigen | -1.20 | 0.033 |
| *Adora2a* | adenosine A2a receptor | -1.57 | 0.004 |
