## Supplementary Table 3. Long-term + Acute DEGs in Males. for "Enduring and sex-specific changes in hippocampal gene expression after a subchronic immune challenge"

**Supplementary Table 3. Differentially expressed genes in the hippocampus of males in response to acute immune challenge, three months after prior subchronic immune challenge (Long-term + Acute condition males).**

| **Gene** | **Gene Name** | **Log2 Fold Change** | **P-value** |
| --- | --- | --- | --- |
| *Isl1* | ISL1 transcription factor, LIM/homeodomain | 3.63 | 0.039 |
| *Prdm12* | PR domain containing 12 | 3.58 | 0.018 |
| *A230065H16Rik* | RIKEN cDNA A230065H16 gene | 2.74 | 0.004 |
| *Gpr151* | G protein-coupled receptor 151 | 2.60 | 0.004 |
| *Eomes* | eomesodermin | 2.24 | 0.004 |
| *Barhl2* | BarH-like 2 (Drosophila) | 2.16 | 0.022 |
| *Gm5741* | predicted gene 5741 | 1.99 | 0.007 |
| *Chrna3* | cholinergic receptor, nicotinic, alpha polypeptide 3 | 1.84 | 0.004 |
| *Slc10a4* | solute carrier family 10 (sodium/bile acid cotransporter family), member 4 | 1.82 | 0.026 |
| *Sstr5* | somatostatin receptor 5 | 1.80 | 0.026 |
| *Chrnb3* | cholinergic receptor, nicotinic, beta polypeptide 3 | 1.78 | 0.004 |
| *Sln* | sarcolipin | 1.75 | 0.031 |
| *Samd3* | sterile alpha motif domain containing 3 | 1.72 | 0.004 |
| *Ptprv* | protein tyrosine phosphatase, receptor type, V | 1.69 | 0.004 |
| *Pcsk9* | proprotein convertase subtilisin/kexin type 9 | 1.61 | 0.004 |
| *Col6a3* | collagen, type VI, alpha 3 | 1.55 | 0.004 |
| *Irs4* | insulin receptor substrate 4 | 1.40 | 0.007 |
| *Baiap3* | BAI1-associated protein 3 | 1.37 | 0.004 |
| *Slc5a7* | solute carrier family 5 (choline transporter), member 7 | 1.36 | 0.004 |
| *Igfbpl1* | insulin-like growth factor binding protein-like 1 | 1.34 | 0.004 |
| *Scn5a* | sodium channel, voltage-gated, type V, alpha | 1.22 | 0.004 |
| *Igfn1* | immunoglobulin-like and fibronectin type III domain containing 1 | 1.17 | 0.004 |
| *Magel2* | melanoma antigen, family L, 2 | 1.15 | 0.004 |
| *Slc35d3* | solute carrier family 35, member D3 | 1.13 | 0.020 |
| *Zic4* | zinc finger protein of the cerebellum 4 | 1.10 | 0.004 |
| *Chrnb4* | cholinergic receptor, nicotinic, beta polypeptide 4 | 1.06 | 0.031 |
| *Tac1* | tachykinin 1 | 1.05 | 0.004 |
| *Pld5* | phospholipase D family, member 5 | 1.05 | 0.004 |
| *Gpr149* | G protein-coupled receptor 149 | 1.03 | 0.044 |
| *Six3* | sine oculis-related homeobox 3 | 0.99 | 0.039 |
| *Penk* | preproenkephalin | 0.96 | 0.004 |
| *Drd2* | dopamine receptor D2 | 0.94 | 0.044 |
| *Arhgap36* | Rho GTPase activating protein 36 | 0.94 | 0.024 |
| *Cartpt* | CART prepropeptide | 0.93 | 0.020 |
| *Ano1* | anoctamin 1, calcium activated chloride channel | 0.93 | 0.004 |
| *Ebf3* | early B cell factor 3 | 0.91 | 0.029 |
| *Ecel1* | endothelin converting enzyme-like 1 | 0.86 | 0.004 |
| *Adora2a* | adenosine A2a receptor | 0.85 | 0.029 |
| *Zic1* | zinc finger protein of the cerebellum 1 | 0.83 | 0.004 |
| *Col6a4* | collagen, type VI, alpha 4 | 0.81 | 0.004 |
| *Foxp2* | forkhead box P2 | 0.81 | 0.015 |
| *Tacr1* | tachykinin receptor 1 | 0.79 | 0.039 |
| *Igsf1* | immunoglobulin superfamily, member 1 | 0.78 | 0.004 |
| *AW551984* | expressed sequence AW551984 | 0.78 | 0.004 |
| *Arhgap6* | Rho GTPase activating protein 6 | 0.77 | 0.004 |
| *Syt6* | synaptotagmin VI | 0.75 | 0.004 |
| *Ano2* | anoctamin 2 | 0.74 | 0.015 |
| *Lrrc55* | leucine rich repeat containing 55 | 0.73 | 0.004 |
| *Zic3* | zinc finger protein of the cerebellum 3 | 0.72 | 0.029 |
| *Drd1a* | dopamine receptor D1 | 0.69 | 0.010 |
| *Zcchc12* | zinc finger, CCHC domain containing 12 | 0.67 | 0.004 |
| *Cacng5* | calcium channel, voltage-dependent, gamma subunit 5 | 0.66 | 0.004 |
| *Susd2* | sushi domain containing 2 | 0.65 | 0.046 |
| *Dlk1* | delta-like 1 homolog (Drosophila) | 0.65 | 0.004 |
| *Gpx3* | glutathione peroxidase 3 | 0.65 | 0.029 |
| *Peg10* | paternally expressed 10 | 0.59 | 0.004 |
| *Lcn2* | lipocalin 2 | -0.74 | 0.010 |
| *Fermt1* | fermitin family member 1 | -0.79 | 0.020 |
