## Supplementary Table 4. Long-term + Acute DEGs in Females. for "Enduring and sex-specific changes in hippocampal gene expression after a subchronic immune challenge"

**Supplementary Table 4. Differentially expressed genes in the hippocampus of females in response to acute immune challenge, three months after prior subchronic immune challenge (Long-term + Acute condition females).**

| **Gene** | **Gene Name** | **Log2 Fold Change** | **P-value** |
| --- | --- | --- | --- |
| *Cga* | glycoprotein hormones, alpha subunit | inf | 0.004 |
| *Serpina9* | serine (or cysteine) peptidase inhibitor, clade A (alpha-1 antiproteinase, antitrypsin), member 9 | 3.86 | 0.004 |
| *Gm10635* | predicted gene 10635 | 3.75 | 0.022 |
| *Cd4* | CD4 antigen | 3.59 | 0.004 |
| *Nxf7* | nuclear RNA export factor 7 | 3.46 | 0.004 |
| *Ntrk1* | neurotrophic tyrosine kinase, receptor, type 1 | 3.10 | 0.007 |
| *Adora2a* | adenosine A2a receptor | 2.95 | 0.004 |
| *4930547N16Rik* | PARP1 binding protein | 2.64 | 0.004 |
| *Ptprv* | protein tyrosine phosphatase, receptor type, V | 2.57 | 0.004 |
| *Chat* | choline acetyltransferase | 2.56 | 0.004 |
| *Drd2* | dopamine receptor D2 | 2.54 | 0.004 |
| *Cd6* | CD6 antigen | 2.46 | 0.004 |
| *Scn4b* | sodium channel, type IV, beta | 2.41 | 0.004 |
| *Gpr6* | G protein-coupled receptor 6 | 2.33 | 0.004 |
| *Ido1* | indoleamine 2,3-dioxygenase 1 | 2.33 | 0.004 |
| *Gpr88* | G-protein coupled receptor 88 | 2.32 | 0.004 |
| *Lhx8* | LIM homeobox protein 8 | 2.32 | 0.044 |
| *Pomc* | pro-opiomelanocortin-alpha | 2.14 | 0.004 |
| *Slc5a7* | solute carrier family 5 (choline transporter), member 7 | 2.13 | 0.004 |
| *Sh3rf2* | SH3 domain containing ring finger 2 | 2.09 | 0.004 |
| *Gm765* | predicted gene 765 | 2.05 | 0.031 |
| *Slc35d3* | solute carrier family 35, member D3 | 1.87 | 0.004 |
| *Krt80* | keratin 80 | 1.84 | 0.004 |
| *Dmkn* | dermokine | 1.81 | 0.039 |
| *Tmem90a* | synapse differentiation inducing 1 like | 1.80 | 0.004 |
| *Trhr2* | thyrotropin releasing hormone receptor 2 | 1.79 | 0.007 |
| *A830009L08Rik* | RIKEN cDNA A830009L08 gene | 1.76 | 0.004 |
| *Prss22* | protease, serine 22 | 1.75 | 0.031 |
| *Olfr78* | olfactory receptor 78 | 1.69 | 0.020 |
| *Chrna3* | cholinergic receptor, nicotinic, alpha polypeptide 3 | 1.69 | 0.004 |
| *Klk6* | kallikrein related-peptidase 6 | 1.68 | 0.004 |
| *Foxp2* | forkhead box P2 | 1.68 | 0.004 |
| *Rgs9* | regulator of G-protein signaling 9 | 1.68 | 0.004 |
| *Stat4* | signal transducer and activator of transcription 4 | 1.59 | 0.007 |
| *Rspo1* | R-spondin 1 | 1.58 | 0.004 |
| *Hfe2* | hemochromatosis type 2 (juvenile) | 1.54 | 0.046 |
| *Gpr151* | G protein-coupled receptor 151 | 1.53 | 0.004 |
| *Oprk1* | opioid receptor, kappa 1 | 1.53 | 0.004 |
| *Car3* | carbonic anhydrase 3 | 1.53 | 0.035 |
| *Htr1d* | 5-hydroxytryptamine (serotonin) receptor 1D | 1.51 | 0.024 |
| *Penk* | preproenkephalin | 1.51 | 0.004 |
| *Nexn* | nexilin | 1.49 | 0.004 |
| *Pou4f1* | POU domain, class 4, transcription factor 1 | 1.47 | 0.013 |
| *Chrnb3* | cholinergic receptor, nicotinic, beta polypeptide 3 | 1.46 | 0.004 |
| *Neu2* | neuraminidase 2 | 1.44 | 0.004 |
| *Myl4* | myosin, light polypeptide 4 | 1.43 | 0.004 |
| *Rpe65* | retinal pigment epithelium 65 | 1.39 | 0.004 |
| *Drd1a* | dopamine receptor D1 | 1.38 | 0.004 |
| *Gkn3* | gastrokine 3 | 1.37 | 0.029 |
| *Col24a1* | collagen, type XXIV, alpha 1 | 1.34 | 0.004 |
| *Cdsn* | corneodesmosin | 1.33 | 0.007 |
| *Sspo* | SCO-spondin | 1.33 | 0.004 |
| *Il12a* | interleukin 12a | 1.33 | 0.042 |
| *Ppp1r1b* | protein phosphatase 1, regulatory (inhibitor) subunit 1B | 1.30 | 0.004 |
| *Pde10a* | phosphodiesterase 10A | 1.26 | 0.004 |
| *Cited4* | Cbp/p300-interacting transactivator, with Glu/Asp-rich carboxy-terminal domain, 4 | 1.26 | 0.004 |
| *Tgm3* | transglutaminase 3, E polypeptide | 1.24 | 0.004 |
| *Kcnh5* | potassium voltage-gated channel, subfamily H (eag-related), member 5 | 1.23 | 0.004 |
| *Adra1b* | adrenergic receptor, alpha 1b | 1.23 | 0.004 |
| *Npc1l1* | NPC1-like 1 | 1.15 | 0.026 |
| *A230001M10Rik* | RIKEN cDNA A230001M10 gene | 1.12 | 0.004 |
| *Tcap* | titin-cap | 1.12 | 0.040 |
| *Kcnh4* | potassium voltage-gated channel, subfamily H (eag-related), member 4 | 1.12 | 0.004 |
| *Dkkl1* | dickkopf-like 1 | 1.11 | 0.004 |
| *Tac1* | tachykinin 1 | 1.10 | 0.004 |
| *Rasd2* | RASD family, member 2 | 1.10 | 0.004 |
| *Gm11549* | predicted gene 11549 | 1.08 | 0.004 |
| *Ccbp2* | atypical chemokine receptor 2 | 1.07 | 0.018 |
| *Syt6* | synaptotagmin VI | 1.06 | 0.004 |
| *Cpne9* | copine family member IX | 1.05 | 0.004 |
| *9330117O12* | RIKEN cDNA 9330117O12 gene | 1.05 | 0.004 |
| *6330527O06Rik* | lysosomal-associated membrane protein family, member 5 | 1.05 | 0.004 |
| *Ninj2* | ninjurin 2 | 1.04 | 0.015 |
| *Rasgrp2* | RAS, guanyl releasing protein 2 | 1.04 | 0.004 |
| *Tspan11* | tetraspanin 11 | 1.04 | 0.022 |
| *Enpp6* | ectonucleotide pyrophosphatase/phosphodiesterase 6 | 1.01 | 0.004 |
| *Sowahb* | sosondowah ankyrin repeat domain family member B | 1.00 | 0.004 |
| *Exph5* | exophilin 5 | 1.00 | 0.004 |
| *Prima1* | proline rich membrane anchor 1 | 0.98 | 0.020 |
| *Rasgef1b* | RasGEF domain family, member 1B | 0.97 | 0.004 |
| *Atp6ap1l* | ATPase, H+ transporting, lysosomal accessory protein 1-like | 0.97 | 0.004 |
| *Opalin* | oligodendrocytic myelin paranodal and inner loop protein | 0.97 | 0.004 |
| *2610034M16Rik* | PDZ and pleckstrin homology domains 1 | 0.97 | 0.013 |
| *Satb2* | special AT-rich sequence binding protein 2 | 0.97 | 0.004 |
| *Cpne5* | copine V | 0.97 | 0.004 |
| *Cckbr* | cholecystokinin B receptor | 0.95 | 0.004 |
| *Gpr133* | adhesion G protein-coupled receptor D1 | 0.95 | 0.007 |
| *Six3* | sine oculis-related homeobox 3 | 0.94 | 0.010 |
| *Kcnab3* | potassium voltage-gated channel, shaker-related subfamily, beta member 3 | 0.94 | 0.004 |
| *Rims3* | regulating synaptic membrane exocytosis 3 | 0.93 | 0.004 |
| *Dpp4* | dipeptidylpeptidase 4 | 0.93 | 0.004 |
| *Mylk3* | myosin light chain kinase 3 | 0.91 | 0.049 |
| *Hr* | hairless | 0.91 | 0.004 |
| *Pde7b* | phosphodiesterase 7B | 0.89 | 0.004 |
| *Ankrd34c* | ankyrin repeat domain 34C | 0.89 | 0.031 |
| *Adam33* | a disintegrin and metallopeptidase domain 33 | 0.88 | 0.004 |
| *Blnk* | B cell linker | 0.88 | 0.004 |
| *Gjb1* | gap junction protein, beta 1 | 0.87 | 0.004 |
| *Cyp39a1* | cytochrome P450, family 39, subfamily a, polypeptide 1 | 0.87 | 0.044 |
| *Arpp19* | cAMP-regulated phosphoprotein 19 | 0.86 | 0.004 |
| *Ovol2* | ovo like zinc finger 2 | 0.86 | 0.033 |
| *Rassf3* | Ras association (RalGDS/AF-6) domain family member 3 | 0.86 | 0.004 |
| *Pdyn* | prodynorphin | 0.85 | 0.007 |
| *Creb5* | cAMP responsive element binding protein 5 | 0.85 | 0.029 |
| *Hs3st2* | heparan sulfate (glucosamine) 3-O-sulfotransferase 2 | 0.84 | 0.004 |
| *6330406I15Rik* | mesenteric estrogen dependent adipogenesis | 0.84 | 0.004 |
| *Kcnh7* | potassium voltage-gated channel, subfamily H (eag-related), member 7 | 0.83 | 0.004 |
| *Gabrd* | gamma-aminobutyric acid (GABA) A receptor, subunit delta | 0.83 | 0.004 |
| *Rarb* | retinoic acid receptor, beta | 0.83 | 0.004 |
| *Ngef* | neuronal guanine nucleotide exchange factor | 0.81 | 0.004 |
| *Rasgef1c* | RasGEF domain family, member 1C | 0.81 | 0.004 |
| *Rgs6* | regulator of G-protein signaling 6 | 0.81 | 0.004 |
| *Rgs4* | regulator of G-protein signaling 4 | 0.80 | 0.004 |
| *Mme* | membrane metallo endopeptidase | 0.80 | 0.004 |
| *Mei1* | meiotic double-stranded break formation protein 1 | 0.80 | 0.022 |
| *St6gal2* | beta galactoside alpha 2,6 sialyltransferase 2 | 0.80 | 0.018 |
| *Lrrc55* | leucine rich repeat containing 55 | 0.79 | 0.004 |
| *Serpinb1a* | serine (or cysteine) peptidase inhibitor, clade B, member 1a | 0.79 | 0.004 |
| *Wnt10a* | wingless-type MMTV integration site family, member 10A | 0.79 | 0.026 |
| *Gjc2* | gap junction protein, gamma 2 | 0.78 | 0.004 |
| *Aspa* | aspartoacylase | 0.77 | 0.004 |
| *Tmem88b* | transmembrane protein 88B | 0.77 | 0.004 |
| *Ddit4l* | DNA-damage-inducible transcript 4-like | 0.77 | 0.004 |
| *Dclk3* | doublecortin-like kinase 3 | 0.76 | 0.004 |
| *Kazald1* | Kazal-type serine peptidase inhibitor domain 1 | 0.76 | 0.040 |
| *Hhip* | Hedgehog-interacting protein | 0.76 | 0.004 |
| *Rnf39* | ring finger protein 39 | 0.76 | 0.037 |
| *Galnt6* | UDP-N-acetyl-alpha-D-galactosamine:polypeptide N-acetylgalactosaminyltransferase 6 | 0.76 | 0.004 |
| *Syt2* | synaptotagmin II | 0.76 | 0.004 |
| *Rorb* | RAR-related orphan receptor beta | 0.75 | 0.004 |
| *Efna5* | ephrin A5 | 0.75 | 0.004 |
| *Ugt8a* | UDP galactosyltransferase 8A | 0.74 | 0.004 |
| *Sytl2* | synaptotagmin-like 2 | 0.73 | 0.004 |
| *Mef2c* | myocyte enhancer factor 2C | 0.73 | 0.004 |
| *Cyp2j12* | cytochrome P450, family 2, subfamily j, polypeptide 12 | 0.73 | 0.039 |
| *Cobl* | cordon-bleu WH2 repeat | 0.73 | 0.004 |
| *Tmem132d* | transmembrane protein 132D | 0.73 | 0.004 |
| *Necab3* | N-terminal EF-hand calcium binding protein 3 | 0.73 | 0.004 |
| *Pde1b* | phosphodiesterase 1B, Ca2+-calmodulin dependent | 0.72 | 0.004 |
| *Cbln2* | cerebellin 2 precursor protein | 0.71 | 0.007 |
| *Rxrg* | retinoid X receptor gamma | 0.71 | 0.042 |
| *Prr16* | proline rich 16 | 0.71 | 0.026 |
| *Krt12* | keratin 12 | 0.71 | 0.004 |
| *Robo3* | roundabout guidance receptor 3 | 0.71 | 0.004 |
| *Plxdc1* | plexin domain containing 1 | 0.71 | 0.004 |
| *Dact2* | dishevelled-binding antagonist of beta-catenin 2 | 0.71 | 0.004 |
| *Gcnt4* | glucosaminyl (N-acetyl) transferase 4, core 2 (beta-1,6-N-acetylglucosaminyltransferase) | 0.71 | 0.010 |
| *Accn4* | acid-sensing (proton-gated) ion channel family member 4 | 0.70 | 0.007 |
| *Stard10* | START domain containing 10 | 0.70 | 0.004 |
| *Gucy1a3* | guanylate cyclase 1, soluble, alpha 3 | 0.69 | 0.004 |
| *Fa2h* | fatty acid 2-hydroxylase | 0.68 | 0.004 |
| *Ipcef1* | interaction protein for cytohesin exchange factors 1 | 0.68 | 0.004 |
| *Sorcs1* | sortilin-related VPS10 domain containing receptor 1 | 0.68 | 0.004 |
| *Prr5l* | proline rich 5 like | 0.68 | 0.004 |
| *1110012J17Rik* | microtubule crosslinking factor 1 | 0.68 | 0.004 |
| *Pak7* | p21 protein (Cdc42/Rac)-activated kinase 7 | 0.68 | 0.004 |
| *Sgpp2* | sphingosine-1-phosphate phosphotase 2 | 0.67 | 0.004 |
| *Ramp3* | receptor (calcitonin) activity modifying protein 3 | 0.67 | 0.010 |
| *C130074G19Rik* | RIKEN cDNA C130074G19 gene | 0.67 | 0.004 |
| *Dach1* | dachshund 1 (Drosophila) | 0.67 | 0.015 |
| *Gpr83* | G protein-coupled receptor 83 | 0.67 | 0.007 |
| *Apln* | apelin | 0.67 | 0.004 |
| *Susd2* | sushi domain containing 2 | 0.67 | 0.015 |
| *Mkx* | mohawk homeobox | 0.67 | 0.004 |
| *Hkdc1* | hexokinase domain containing 1 | 0.67 | 0.039 |
| *Epha8* | Eph receptor A8 | 0.66 | 0.026 |
| *Inf2* | inverted formin, FH2 and WH2 domain containing | 0.66 | 0.004 |
| *Mcam* | melanoma cell adhesion molecule | 0.66 | 0.004 |
| *Cabp1* | calcium binding protein 1 | 0.65 | 0.004 |
| *Pllp* | plasma membrane proteolipid | 0.65 | 0.004 |
| *Lrrc10b* | leucine rich repeat containing 10B | 0.64 | 0.004 |
| *Ppp1r14a* | protein phosphatase 1, regulatory (inhibitor) subunit 14A | 0.64 | 0.018 |
| *Igfbp6* | insulin-like growth factor binding protein 6 | 0.63 | 0.007 |
| *Bcas1* | breast carcinoma amplified sequence 1 | 0.63 | 0.004 |
| *Pamr1* | peptidase domain containing associated with muscle regeneration 1 | 0.62 | 0.004 |
| *Tmem125* | transmembrane protein 125 | 0.62 | 0.020 |
| *Stard8* | START domain containing 8 | 0.62 | 0.007 |
| *Pou3f2* | POU domain, class 3, transcription factor 2 | 0.62 | 0.007 |
| *Plekhh1* | pleckstrin homology domain containing, family H (with MyTH4 domain) member 1 | 0.62 | 0.004 |
| *Kcns1* | K+ voltage-gated channel, subfamily S, 1 | 0.62 | 0.007 |
| *Grip2* | glutamate receptor interacting protein 2 | 0.62 | 0.004 |
| *Igsf9b* | immunoglobulin superfamily, member 9B | 0.62 | 0.046 |
| *Kcnk13* | potassium channel, subfamily K, member 13 | 0.61 | 0.033 |
| *Cux2* | cut-like homeobox 2 | 0.61 | 0.004 |
| *Mag* | myelin-associated glycoprotein | 0.61 | 0.004 |
| *Ermn* | ermin, ERM-like protein | 0.61 | 0.004 |
| *Sox2ot* | SOX2 overlapping transcript (non-protein coding) | 0.60 | 0.018 |
| *Camk2n1* | calcium/calmodulin-dependent protein kinase II inhibitor 1 | 0.60 | 0.010 |
| *Prr18* | proline rich 18 | 0.60 | 0.004 |
| *Gpr37* | G protein-coupled receptor 37 | 0.59 | 0.004 |
| *Pvalb* | parvalbumin | 0.59 | 0.004 |
| *Sept4* | septin 4 | 0.59 | 0.004 |
| *Apaf1* | apoptotic peptidase activating factor 1 | -0.59 | 0.004 |
| *Fkbp5* | FK506 binding protein 5 | -0.59 | 0.004 |
| *6030405A18Rik* | serine rich and transmembrane domain containing 1 | -0.59 | 0.004 |
| *Gpc4* | glypican 4 | -0.59 | 0.004 |
| *Xdh* | xanthine dehydrogenase | -0.59 | 0.004 |
| *Emb* | embigin | -0.60 | 0.004 |
| *Rbp1* | retinol binding protein 1, cellular | -0.60 | 0.013 |
| *Nov* | nephroblastoma overexpressed gene | -0.61 | 0.004 |
| *Phactr2* | phosphatase and actin regulator 2 | -0.61 | 0.004 |
| *Islr* | immunoglobulin superfamily containing leucine-rich repeat | -0.61 | 0.007 |
| *AW551984* | expressed sequence AW551984 | -0.62 | 0.031 |
| *Aspg* | asparaginase | -0.62 | 0.042 |
| *Arhgap6* | Rho GTPase activating protein 6 | -0.62 | 0.018 |
| *Crym* | crystallin, mu | -0.62 | 0.004 |
| *Pglyrp1* | peptidoglycan recognition protein 1 | -0.63 | 0.026 |
| *Moxd1* | monooxygenase, DBH-like 1 | -0.63 | 0.029 |
| *Slc6a13* | solute carrier family 6 (neurotransmitter transporter, GABA), member 13 | -0.63 | 0.013 |
| *Serpina3n* | serine (or cysteine) peptidase inhibitor, clade A, member 3N | -0.64 | 0.004 |
| *Lgals9* | lectin, galactose binding, soluble 9 | -0.64 | 0.015 |
| *Slc4a2* | solute carrier family 4 (anion exchanger), member 2 | -0.64 | 0.004 |
| *Mt2* | metallothionein 2 | -0.65 | 0.004 |
| *Nxn* | nucleoredoxin | -0.65 | 0.004 |
| *B4galt1* | UDP-Gal:betaGlcNAc beta 1,4- galactosyltransferase, polypeptide 1 | -0.65 | 0.018 |
| *Cab39l* | calcium binding protein 39-like | -0.65 | 0.004 |
| *Gbp4* | guanylate binding protein 4 | -0.66 | 0.022 |
| *Pgcp* | carboxypeptidase Q | -0.66 | 0.004 |
| *Gjb2* | gap junction protein, beta 2 | -0.67 | 0.004 |
| *Lgals1* | lectin, galactose binding, soluble 1 | -0.67 | 0.007 |
| *Spink8* | serine peptidase inhibitor, Kazal type 8 | -0.67 | 0.018 |
| *Col23a1* | collagen, type XXIII, alpha 1 | -0.67 | 0.004 |
| *Slc29a4* | solute carrier family 29 (nucleoside transporters), member 4 | -0.67 | 0.004 |
| *Vat1l* | vesicle amine transport protein 1 like | -0.67 | 0.004 |
| *Slc9a2* | solute carrier family 9 (sodium/hydrogen exchanger), member 2 | -0.67 | 0.004 |
| *Cgnl1* | cingulin-like 1 | -0.68 | 0.004 |
| *Cdkn1c* | cyclin-dependent kinase inhibitor 1C (P57) | -0.68 | 0.004 |
| *Oasl2* | 2'-5' oligoadenylate synthetase-like 2 | -0.68 | 0.026 |
| *Slc9a4* | solute carrier family 9 (sodium/hydrogen exchanger), member 4 | -0.68 | 0.007 |
| *Nnat* | neuronatin | -0.69 | 0.010 |
| *Osmr* | oncostatin M receptor | -0.69 | 0.007 |
| *Tgfbi* | transforming growth factor, beta induced | -0.69 | 0.013 |
| *Bgn* | biglycan | -0.69 | 0.004 |
| *Angpt1* | angiopoietin 1 | -0.70 | 0.013 |
| *Vcam1* | vascular cell adhesion molecule 1 | -0.71 | 0.004 |
| *Map3k6* | mitogen-activated protein kinase kinase kinase 6 | -0.71 | 0.010 |
| *Hif3a* | hypoxia inducible factor 3, alpha subunit | -0.71 | 0.004 |
| *Sult1a1* | sulfotransferase family 1A, phenol-preferring, member 1 | -0.71 | 0.004 |
| *Oxtr* | oxytocin receptor | -0.71 | 0.004 |
| *Lepr* | leptin receptor | -0.71 | 0.007 |
| *Cp* | ceruloplasmin | -0.71 | 0.010 |
| *Gpr161* | G protein-coupled receptor 161 | -0.71 | 0.020 |
| *Mid1* | midline 1 | -0.72 | 0.004 |
| *Ucp2* | uncoupling protein 2 (mitochondrial, proton carrier) | -0.72 | 0.004 |
| *Pdgfd* | platelet-derived growth factor, D polypeptide | -0.73 | 0.049 |
| *Mrc1* | mannose receptor, C type 1 | -0.73 | 0.010 |
| *Crispld2* | cysteine-rich secretory protein LCCL domain containing 2 | -0.73 | 0.033 |
| *Slc16a9* | solute carrier family 16 (monocarboxylic acid transporters), member 9 | -0.74 | 0.004 |
| *Fibcd1* | fibrinogen C domain containing 1 | -0.74 | 0.004 |
| *Fhad1* | forkhead-associated (FHA) phosphopeptide binding domain 1 | -0.74 | 0.004 |
| *Nr2f2* | nuclear receptor subfamily 2, group F, member 2 | -0.74 | 0.004 |
| *Fmod* | fibromodulin | -0.74 | 0.004 |
| *Frem1* | Fras1 related extracellular matrix protein 1 | -0.75 | 0.004 |
| *A430107O13Rik* | cadherin-like and PC-esterase domain containing 1 | -0.75 | 0.020 |
| *Igfbp3* | insulin-like growth factor binding protein 3 | -0.75 | 0.004 |
| *Emp1* | epithelial membrane protein 1 | -0.75 | 0.037 |
| *Col18a1* | collagen, type XVIII, alpha 1 | -0.76 | 0.004 |
| *Eln* | elastin | -0.76 | 0.004 |
| *Fbln1* | fibulin 1 | -0.77 | 0.004 |
| *Slc31a1* | solute carrier family 31, member 1 | -0.77 | 0.004 |
| *Ptpn14* | protein tyrosine phosphatase, non-receptor type 14 | -0.78 | 0.004 |
| *Cpne7* | copine VII | -0.79 | 0.004 |
| *F13a1* | coagulation factor XIII, A1 subunit | -0.79 | 0.020 |
| *Lgals3bp* | lectin, galactoside-binding, soluble, 3 binding protein | -0.80 | 0.004 |
| *Bmp6* | bone morphogenetic protein 6 | -0.80 | 0.004 |
| *Ifitm3* | interferon induced transmembrane protein 3 | -0.81 | 0.004 |
| *Myof* | myoferlin | -0.82 | 0.004 |
| *Serping1* | serine (or cysteine) peptidase inhibitor, clade G, member 1 | -0.84 | 0.004 |
| *Mdfic* | MyoD family inhibitor domain containing | -0.84 | 0.004 |
| *Fzd4* | frizzled class receptor 4 | -0.84 | 0.004 |
| *Spint2* | serine protease inhibitor, Kunitz type 2 | -0.84 | 0.004 |
| *Trhr* | thyrotropin releasing hormone receptor | -0.85 | 0.004 |
| *Tinagl1* | tubulointerstitial nephritis antigen-like 1 | -0.85 | 0.004 |
| *Bmp7* | bone morphogenetic protein 7 | -0.86 | 0.004 |
| *Gpx3* | glutathione peroxidase 3 | -0.86 | 0.004 |
| *Gstm2* | glutathione S-transferase, mu 2 | -0.86 | 0.026 |
| *Stra6* | stimulated by retinoic acid gene 6 | -0.87 | 0.004 |
| *Slc37a2* | solute carrier family 37 (glycerol-3-phosphate transporter), member 2 | -0.87 | 0.004 |
| *Cybrd1* | cytochrome b reductase 1 | -0.87 | 0.007 |
| *Ppl* | periplakin | -0.87 | 0.004 |
| *Eya2* | EYA transcriptional coactivator and phosphatase 2 | -0.89 | 0.037 |
| *Ptgds* | prostaglandin D2 synthase (brain) | -0.89 | 0.004 |
| *Pla2g5* | phospholipase A2, group V | -0.89 | 0.039 |
| *Npr3* | natriuretic peptide receptor 3 | -0.89 | 0.004 |
| *C5ar1* | complement component 5a receptor 1 | -0.90 | 0.031 |
| *Cox6b2* | cytochrome c oxidase subunit VIb polypeptide 2 | -0.90 | 0.022 |
| *Slc16a12* | solute carrier family 16 (monocarboxylic acid transporters), member 12 | -0.91 | 0.007 |
| *Olfr1033* | olfactory receptor 1033 | -0.91 | 0.049 |
| *Prph* | peripherin | -0.91 | 0.029 |
| *Cd59a* | CD59a antigen | -0.92 | 0.004 |
| *Zfp185* | zinc finger protein 185 | -0.92 | 0.004 |
| *Cpxm2* | carboxypeptidase X 2 (M14 family) | -0.93 | 0.018 |
| *Thbs1* | thrombospondin 1 | -0.93 | 0.010 |
| *Perp* | PERP, TP53 apoptosis effector | -0.93 | 0.004 |
| *Itih2* | inter-alpha trypsin inhibitor, heavy chain 2 | -0.93 | 0.004 |
| *Ppp1r3b* | protein phosphatase 1, regulatory (inhibitor) subunit 3B | -0.95 | 0.024 |
| *Aox3* | aldehyde oxidase 3 | -0.95 | 0.004 |
| *Llgl2* | lethal giant larvae homolog 2 (Drosophila) | -0.96 | 0.031 |
| *St6galnac2* | ST6 (alpha-N-acetyl-neuraminyl-2,3-beta-galactosyl-1,3)-N-acetylgalactosaminide alpha-2,6-sialyltransferase 2 | -0.96 | 0.004 |
| *Gbp2* | guanylate binding protein 2 | -0.97 | 0.004 |
| *Inmt* | indolethylamine N-methyltransferase | -0.98 | 0.049 |
| *Irf7* | interferon regulatory factor 7 | -0.98 | 0.004 |
| *Il2rg* | interleukin 2 receptor, gamma chain | -0.98 | 0.026 |
| *Lyve1* | lymphatic vessel endothelial hyaluronan receptor 1 | -0.98 | 0.013 |
| *Mgp* | matrix Gla protein | -0.99 | 0.004 |
| *Gpx8* | glutathione peroxidase 8 (putative) | -1.00 | 0.004 |
| *Aldh1a2* | aldehyde dehydrogenase family 1, subfamily A2 | -1.00 | 0.004 |
| *Car12* | carbonic anhydrase 12 | -1.03 | 0.004 |
| *Mc4r* | melanocortin 4 receptor | -1.03 | 0.007 |
| *Spp1* | secreted phosphoprotein 1 | -1.03 | 0.004 |
| *Gpr182* | G protein-coupled receptor 182 | -1.04 | 0.004 |
| *Sulf1* | sulfatase 1 | -1.05 | 0.004 |
| *H2-Eb1* | histocompatibility 2, class II antigen E beta | -1.05 | 0.010 |
| *A130040M12Rik* | RIKEN cDNA A130040M12 gene | -1.05 | 0.004 |
| *Ptgis* | prostaglandin I2 (prostacyclin) synthase | -1.06 | 0.035 |
| *Cyp1b1* | cytochrome P450, family 1, subfamily b, polypeptide 1 | -1.06 | 0.004 |
| *Prrg4* | proline rich Gla (G-carboxyglutamic acid) 4 (transmembrane) | -1.07 | 0.010 |
| *Sgms2* | sphingomyelin synthase 2 | -1.08 | 0.004 |
| *Wfikkn2* | WAP, follistatin/kazal, immunoglobulin, kunitz and netrin domain containing 2 | -1.08 | 0.004 |
| *Bst2* | bone marrow stromal cell antigen 2 | -1.09 | 0.004 |
| *Gpr101* | G protein-coupled receptor 101 | -1.09 | 0.004 |
| *Icam1* | intercellular adhesion molecule 1 | -1.10 | 0.004 |
| *Trpm3* | transient receptor potential cation channel, subfamily M, member 3 | -1.11 | 0.004 |
| *Ltc4s* | leukotriene C4 synthase | -1.11 | 0.004 |
| *Best3* | bestrophin 3 | -1.13 | 0.024 |
| *Steap2* | six transmembrane epithelial antigen of prostate 2 | -1.13 | 0.004 |
| *Slc6a12* | solute carrier family 6 (neurotransmitter transporter, betaine/GABA), member 12 | -1.14 | 0.004 |
| *Cd74* | CD74 antigen (invariant polypeptide of major histocompatibility complex, class II antigen-associated) | -1.15 | 0.004 |
| *Ms4a6d* | membrane-spanning 4-domains, subfamily A, member 6D | -1.15 | 0.004 |
| *Pcolce* | procollagen C-endopeptidase enhancer protein | -1.16 | 0.004 |
| *Fam115c* | TRPM8 channel-associated factor 2 | -1.17 | 0.022 |
| *Olfm4* | olfactomedin 4 | -1.19 | 0.004 |
| *Ifi44* | interferon-induced protein 44 | -1.20 | 0.004 |
| *Epn3* | epsin 3 | -1.20 | 0.004 |
| *Glb1l2* | galactosidase, beta 1-like 2 | -1.21 | 0.004 |
| *Rtp1* | receptor transporter protein 1 | -1.22 | 0.007 |
| *H2-Ab1* | histocompatibility 2, class II antigen A, beta 1 | -1.22 | 0.004 |
| *Itpripl1* | inositol 1,4,5-triphosphate receptor interacting protein-like 1 | -1.22 | 0.004 |
| *Slc6a20a* | solute carrier family 6 (neurotransmitter transporter), member 20A | -1.23 | 0.004 |
| *B4galnt3* | beta-1,4-N-acetyl-galactosaminyl transferase 3 | -1.24 | 0.029 |
| *Msx1as* | msh homeobox 1 opposite strand | -1.24 | 0.004 |
| *Trh* | thyrotropin releasing hormone | -1.27 | 0.007 |
| *H2-Aa* | histocompatibility 2, class II antigen A, alpha | -1.28 | 0.004 |
| *Crhr2* | corticotropin releasing hormone receptor 2 | -1.28 | 0.004 |
| *Cdh1* | cadherin 1 | -1.28 | 0.004 |
| *Col9a3* | collagen, type IX, alpha 3 | -1.28 | 0.004 |
| *Dcn* | decorin | -1.29 | 0.004 |
| *Nmbr* | neuromedin B receptor | -1.29 | 0.004 |
| *D14Ertd668e* | PHD finger protein 11D | -1.30 | 0.033 |
| *Siglec1* | sialic acid binding Ig-like lectin 1, sialoadhesin | -1.32 | 0.007 |
| *Ifitm1* | interferon induced transmembrane protein 1 | -1.36 | 0.004 |
| *Rab20* | RAB20, member RAS oncogene family | -1.37 | 0.026 |
| *Tgtp1* | T cell specific GTPase 1 | -1.39 | 0.040 |
| *Pon3* | paraoxonase 3 | -1.40 | 0.004 |
| *Sostdc1* | sclerostin domain containing 1 | -1.41 | 0.004 |
| *Dlk1* | delta-like 1 homolog (Drosophila) | -1.41 | 0.004 |
| *Lcn2* | lipocalin 2 | -1.44 | 0.004 |
| *H2-Q1* | histocompatibility 2, Q region locus 1 | -1.45 | 0.046 |
| *Sfrp5* | secreted frizzled-related sequence protein 5 | -1.45 | 0.004 |
| *Mc3r* | melanocortin 3 receptor | -1.46 | 0.048 |
| *Lbp* | lipopolysaccharide binding protein | -1.46 | 0.004 |
| *Lefty2* | left-right determination factor 2 | -1.46 | 0.024 |
| *Tuba1c* | tubulin, alpha 1C | -1.46 | 0.004 |
| *Col17a1* | collagen, type XVII, alpha 1 | -1.46 | 0.004 |
| *Ccdc135* | dynein regulatory complex subunit 7 | -1.47 | 0.004 |
| *Igfbp2* | insulin-like growth factor binding protein 2 | -1.47 | 0.004 |
| *Rdh5* | retinol dehydrogenase 5 | -1.47 | 0.004 |
| *Krt8* | keratin 8 | -1.48 | 0.004 |
| *Lmx1a* | LIM homeobox transcription factor 1 alpha | -1.48 | 0.004 |
| *Ch25h* | cholesterol 25-hydroxylase | -1.49 | 0.004 |
| *Slc13a4* | solute carrier family 13 (sodium/sulfate symporters), member 4 | -1.50 | 0.004 |
| *Slc39a4* | solute carrier family 39 (zinc transporter), member 4 | -1.51 | 0.004 |
| *Rbm47* | RNA binding motif protein 47 | -1.51 | 0.004 |
| *Mpzl2* | myelin protein zero-like 2 | -1.52 | 0.004 |
| *Abca4* | ATP-binding cassette, sub-family A (ABC1), member 4 | -1.52 | 0.004 |
| *Plek2* | pleckstrin 2 | -1.55 | 0.010 |
| *Enpp2* | ectonucleotide pyrophosphatase/phosphodiesterase 2 | -1.66 | 0.004 |
| *Cldn1* | claudin 1 | -1.66 | 0.004 |
| *Krt18* | keratin 18 | -1.67 | 0.004 |
| *Slco1a5* | solute carrier organic anion transporter family, member 1a5 | -1.67 | 0.018 |
| *Igf2* | insulin-like growth factor 2 | -1.69 | 0.004 |
| *Col8a2* | collagen, type VIII, alpha 2 | -1.71 | 0.004 |
| *Ace* | angiotensin I converting enzyme (peptidyl-dipeptidase A) 1 | -1.72 | 0.004 |
| *Il31ra* | interleukin 31 receptor A | -1.77 | 0.013 |
| *Sema3b* | sema domain, immunoglobulin domain (Ig), short basic domain, secreted, (semaphorin) 3B | -1.78 | 0.004 |
| *1110059M19Rik* | proline rich 32 | -1.79 | 0.004 |
| *Dmrt3* | doublesex and mab-3 related transcription factor 3 | -1.80 | 0.022 |
| *Steap4* | STEAP family member 4 | -1.81 | 0.004 |
| *Gyltl1b* | glycosyltransferase-like 1B | -1.81 | 0.007 |
| *Cdh3* | cadherin 3 | -1.83 | 0.004 |
| *Otx2* | orthodenticle homeobox 2 | -1.85 | 0.004 |
| *2810433D01Rik* | RIKEN cDNA 2810433D01 gene | -1.86 | 0.040 |
| *Scara5* | scavenger receptor class A, member 5 | -1.86 | 0.004 |
| *Dsc3* | desmocollin 3 | -1.87 | 0.004 |
| *Col4a3* | collagen, type IV, alpha 3 | -1.89 | 0.004 |
| *Wdr72* | WD repeat domain 72 | -1.90 | 0.004 |
| *Adh1* | alcohol dehydrogenase 1 (class I) | -1.90 | 0.007 |
| *Il17re* | interleukin 17 receptor E | -1.94 | 0.004 |
| *Slfn4* | schlafen 4 | -1.98 | 0.004 |
| *Plac8* | placenta-specific 8 | -1.98 | 0.004 |
| *Trpv4* | transient receptor potential cation channel, subfamily V, member 4 | -2.00 | 0.004 |
| *Col8a1* | collagen, type VIII, alpha 1 | -2.02 | 0.004 |
| *Kl* | klotho | -2.05 | 0.004 |
| *F5* | coagulation factor V | -2.08 | 0.004 |
| *Clic6* | chloride intracellular channel 6 | -2.09 | 0.004 |
| *Tmem184a* | transmembrane protein 184a | -2.11 | 0.004 |
| *Col4a4* | collagen, type IV, alpha 4 | -2.17 | 0.004 |
| *Kcne2* | potassium voltage-gated channel, Isk-related subfamily, gene 2 | -2.23 | 0.004 |
| *A2m* | alpha-2-macroglobulin | -2.24 | 0.004 |
| *Wdr86* | WD repeat domain 86 | -2.26 | 0.004 |
| *1500015O10Rik* | RIKEN cDNA 1500015O10 gene | -2.27 | 0.004 |
| *Prlr* | prolactin receptor | -2.30 | 0.004 |
| *Tmem27* | transmembrane protein 27 | -2.32 | 0.004 |
| *Slc16a8* | solute carrier family 16 (monocarboxylic acid transporters), member 8 | -2.41 | 0.004 |
| *Aqp1* | aquaporin 1 | -2.42 | 0.004 |
| *Bhmt* | betaine-homocysteine methyltransferase | -2.43 | 0.035 |
| *Wfdc2* | WAP four-disulfide core domain 2 | -2.50 | 0.004 |
| *Tmem72* | transmembrane protein 72 | -2.51 | 0.004 |
| *Cldn9* | claudin 9 | -2.52 | 0.013 |
| *Steap1* | six transmembrane epithelial antigen of the prostate 1 | -2.53 | 0.004 |
| *Tc2n* | tandem C2 domains, nuclear | -2.57 | 0.004 |
| *Sult1c1* | sulfotransferase family, cytosolic, 1C, member 1 | -2.60 | 0.013 |
| *Ttr* | transthyretin | -2.62 | 0.004 |
| *Folr1* | folate receptor 1 (adult) | -2.62 | 0.004 |
| *Slc4a5* | solute carrier family 4, sodium bicarbonate cotransporter, member 5 | -2.63 | 0.004 |
| *Rrh* | retinal pigment epithelium derived rhodopsin homolog | -2.63 | 0.013 |
| *Sult1c2* | sulfotransferase family, cytosolic, 1C, member 2 | -2.64 | 0.004 |
| *Tmprss11a* | transmembrane protease, serine 11a | -2.83 | 0.004 |
| *Cldn2* | claudin 2 | -2.94 | 0.004 |
| *Oca2* | oculocutaneous albinism II | -3.00 | 0.004 |
| *Sele* | selectin, endothelial cell | -3.18 | 0.013 |
| *Dpep1* | dipeptidase 1 (renal) | -3.24 | 0.004 |
