## Supplementary Table 5. Long-term + Acute shared between Males.and Females. for "Enduring and sex-specific changes in hippocampal gene expression after a subchronic immune challenge"

**Supplementary Table 5. Commonly dysregulated genes in the hippocampus of males and females in the Long-term+Acute condition.**

|  | | **Male** | | **Female** | |
| --- | --- | --- | --- | --- | --- |
| **Gene** | **Gene Name** | **Log2 Fold Change** | **P-value** | **Log2 Fold Change** | **P-value** |
| *Adora2a* | adenosine A2a receptor | 0.85 | 0.029 | 2.95 | 0.004 |
| *Arhgap6* | Rho GTPase activating protein 6 | 0.77 | 0.004 | -0.62 | 0.018 |
| *AW551984* | expressed sequence AW551984 | 0.78 | 0.004 | -0.62 | 0.031 |
| *Chrna3* | cholinergic receptor, nicotinic, alpha polypeptide 3 | 1.84 | 0.004 | 1.69 | 0.004 |
| *Chrnb3* | cholinergic receptor, nicotinic, beta polypeptide 3 | 1.78 | 0.004 | 1.46 | 0.004 |
| *Dlk1* | delta-like 1 homolog (Drosophila) | 0.65 | 0.004 | -1.41 | 0.004 |
| *Drd1a* | dopamine receptor D1 | 0.69 | 0.010 | 1.38 | 0.004 |
| *Drd2* | dopamine receptor D2 | 0.94 | 0.044 | 2.54 | 0.004 |
| *Gpr151* | G protein-coupled receptor 151 | 2.60 | 0.004 | 1.53 | 0.004 |
| *Gpx3* | glutathione peroxidase 3 | 0.65 | 0.029 | -0.86 | 0.004 |
| *Foxp2* | forkhead box P2 | 0.81 | 0.015 | 1.68 | 0.004 |
| *Lcn2* | lipocalin 2 | -0.74 | 0.010 | -1.44 | 0.004 |
| *Lrrc55* | leucine rich repeat containing 55 | 0.73 | 0.004 | 0.79 | 0.004 |
| *Penk* | preproenkephalin | 0.96 | 0.004 | 1.51 | 0.004 |
| *Ptprv* | protein tyrosine phosphatase, receptor type, V | 1.69 | 0.004 | 2.57 | 0.004 |
| *Six3* | sine oculis-related homeobox 3 | 0.99 | 0.039 | 0.94 | 0.010 |
| *Slc5a7* | solute carrier family 5 (choline transporter), member 7 | 1.36 | 0.004 | 2.13 | 0.004 |
| *Slc35d3* | solute carrier family 35, member D3 | 1.13 | 0.020 | 1.87 | 0.004 |
| *Susd2* | sushi domain containing 2 | 0.65 | 0.046 | 0.67 | 0.015 |
| *Syt6* | synaptotagmin VI | 0.75 | 0.004 | 1.06 | 0.004 |
| *Tac1* | tachykinin 1 | 1.05 | 0.004 | 1.10 | 0.004 |
