## Supplementary Table 6. Male and Female Biased Genes. for "Enduring and sex-specific changes in hippocampal gene expression after a subchronic immune challenge"

**Supplementary Table 6. Genes higher in the female hippocampus at baseline (“female-biased” genes; positive Log2 Fold Change) and genes higher in the male hippocampus at baseline (“male-biased” genes; negative Log2 Fold Change).**

| **Genes higher in females at baseline** | | | |
| --- | --- | --- | --- |
| **Gene** | **Gene Name** | **Log2 Fold Change** | **P-value** |
| *Xist* | inactive X specific transcripts | 8.28 | 0.004 |
| *H2-Q6* | histocompatibility 2, Q region locus 6 | 2.57 | 0.007 |
| *Prdm6* | PR domain containing 6 | 2.48 | 0.010 |
| *Slc6a12* | solute carrier family 6 (neurotransmitter transporter, betaine/GABA), member 12 | 2.44 | 0.004 |
| *Mgl2* | macrophage galactose N-acetyl-galactosamine specific lectin 2 | 2.08 | 0.004 |
| *Slc22a2* | solute carrier family 22 (organic cation transporter), member 2 | 2.06 | 0.004 |
| *Fgfbp1* | fibroblast growth factor binding protein 1 | 1.89 | 0.004 |
| *Cdh1* | cadherin 1 | 1.71 | 0.004 |
| *Spp1* | secreted phosphoprotein 1 | 1.71 | 0.004 |
| *Wnt6* | wingless-type MMTV integration site family, member 6 | 1.70 | 0.004 |
| *Foxc2* | forkhead box C2 | 1.69 | 0.004 |
| *Fam180a* | family with sequence similarity 180, member A | 1.65 | 0.004 |
| *Slc47a1* | solute carrier family 47, member 1 | 1.62 | 0.004 |
| *Ptgdr* | prostaglandin D receptor | 1.51 | 0.004 |
| *Fmod* | fibromodulin | 1.50 | 0.004 |
| *H2-Q1* | histocompatibility 2, Q region locus 1 | 1.50 | 0.004 |
| *Gbp4* | guanylate binding protein 4 | 1.47 | 0.004 |
| *Slc26a7* | solute carrier family 26, member 7 | 1.46 | 0.035 |
| *Col3a1* | collagen, type III, alpha 1 | 1.45 | 0.004 |
| *Sphk1* | sphingosine kinase 1 | 1.40 | 0.004 |
| *Ptgds* | prostaglandin D2 synthase (brain) | 1.40 | 0.004 |
| *Ptgfr* | prostaglandin F receptor | 1.39 | 0.007 |
| *Crabp2* | cellular retinoic acid binding protein II | 1.38 | 0.004 |
| *Col1a1* | collagen, type I, alpha 1 | 1.38 | 0.004 |
| *Cd4* | CD4 antigen | 1.38 | 0.018 |
| *Col6a4* | collagen, type VI, alpha 4 | 1.36 | 0.004 |
| *Ifi47* | interferon gamma inducible protein 47 | 1.36 | 0.031 |
| *Slc6a13* | solute carrier family 6 (neurotransmitter transporter, GABA), member 13 | 1.34 | 0.004 |
| *Slc22a6* | solute carrier family 22 (organic anion transporter), member 6 | 1.33 | 0.004 |
| *Asgr1* | asialoglycoprotein receptor 1 | 1.33 | 0.004 |
| *Ifi44* | interferon-induced protein 44 | 1.30 | 0.010 |
| *Aldh1a2* | aldehyde dehydrogenase family 1, subfamily A2 | 1.30 | 0.004 |
| *Gpr81* | hydrocarboxylic acid receptor 1 | 1.28 | 0.007 |
| *C2* | complement component 2 (within H-2S) | 1.26 | 0.004 |
| *Foxd1* | forkhead box D1 | 1.23 | 0.004 |
| *Mrgprf* | MAS-related GPR, member F | 1.23 | 0.013 |
| *Adamtsl3* | ADAMTS-like 3 | 1.20 | 0.004 |
| *H2-Ab1* | histocompatibility 2, class II antigen A, beta 1 | 1.20 | 0.004 |
| *Drd2* | dopamine receptor D2 | 1.17 | 0.004 |
| *Igfbp6* | insulin-like growth factor binding protein 6 | 1.17 | 0.004 |
| *Itih2* | inter-alpha trypsin inhibitor, heavy chain 2 | 1.13 | 0.004 |
| *Mpzl2* | myelin protein zero-like 2 | 1.12 | 0.004 |
| *Aox3* | aldehyde oxidase 3 | 1.11 | 0.004 |
| *Gjb2* | gap junction protein, beta 2 | 1.10 | 0.004 |
| *Cd74* | CD74 antigen (invariant polypeptide of major histocompatibility complex, class II antigen-associated) | 1.10 | 0.004 |
| *Scn4b* | sodium channel, type IV, beta | 1.10 | 0.004 |
| *Gbp2* | guanylate binding protein 2 | 1.10 | 0.004 |
| *Col1a2* | collagen, type I, alpha 2 | 1.07 | 0.004 |
| *Islr* | immunoglobulin superfamily containing leucine-rich repeat | 1.05 | 0.004 |
| *Slc13a4* | solute carrier family 13 (sodium/sulfate symporters), member 4 | 1.04 | 0.004 |
| *H2-Aa* | histocompatibility 2, class II antigen A, alpha | 1.01 | 0.020 |
| *Aebp1* | AE binding protein 1 | 1.01 | 0.004 |
| *Slc13a3* | solute carrier family 13 (sodium-dependent dicarboxylate transporter), member 3 | 0.99 | 0.004 |
| *1700030C10Rik* | RIKEN cDNA 1700030C10 gene | 0.99 | 0.024 |
| *H2-T23* | histocompatibility 2, T region locus 23 | 0.97 | 0.004 |
| *Sned1* | sushi, nidogen and EGF-like domains 1 | 0.97 | 0.004 |
| *Gpr88* | G-protein coupled receptor 88 | 0.97 | 0.004 |
| *Isg15* | ISG15 ubiquitin-like modifier | 0.95 | 0.044 |
| *Ifit1* | interferon-induced protein with tetratricopeptide repeats 1 | 0.94 | 0.010 |
| *Col4a6* | collagen, type IV, alpha 6 | 0.94 | 0.024 |
| *Foxp2* | forkhead box P2 | 0.94 | 0.004 |
| *Serping1* | serine (or cysteine) peptidase inhibitor, clade G, member 1 | 0.94 | 0.004 |
| *H2-Q4* | histocompatibility 2, Q region locus 4 | 0.93 | 0.004 |
| *Mrc1* | mannose receptor, C type 1 | 0.93 | 0.004 |
| *Efemp1* | epidermal growth factor-containing fibulin-like extracellular matrix protein 1 | 0.93 | 0.004 |
| *Nat1* | N-acetyl transferase 1 | 0.93 | 0.035 |
| *Anpep* | alanyl (membrane) aminopeptidase | 0.92 | 0.004 |
| *Bmp7* | bone morphogenetic protein 7 | 0.89 | 0.004 |
| *Ogn* | osteoglycin | 0.89 | 0.004 |
| *Dpp4* | dipeptidylpeptidase 4 | 0.88 | 0.018 |
| *Gbp3* | guanylate binding protein 3 | 0.88 | 0.004 |
| *Oasl2* | 2'-5' oligoadenylate synthetase-like 2 | 0.88 | 0.004 |
| *Osr1* | odd-skipped related 1 (Drosophila) | 0.87 | 0.015 |
| *Drd1a* | dopamine receptor D1 | 0.87 | 0.004 |
| *Pcolce* | procollagen C-endopeptidase enhancer protein | 0.86 | 0.004 |
| *Adora2a* | adenosine A2a receptor | 0.86 | 0.010 |
| *Wnt10a* | wingless-type MMTV integration site family, member 10A | 0.84 | 0.026 |
| *Col9a2* | collagen, type IX, alpha 2 | 0.82 | 0.004 |
| *Nupr1* | nuclear protein transcription regulator 1 | 0.82 | 0.018 |
| *Rasgef1c* | RasGEF domain family, member 1C | 0.81 | 0.004 |
| *Lamc2* | laminin, gamma 2 | 0.81 | 0.004 |
| *Emilin1* | elastin microfibril interfacer 1 | 0.80 | 0.042 |
| *Capn11* | calpain 11 | 0.79 | 0.020 |
| *Sema3a* | sema domain, immunoglobulin domain (Ig), short basic domain, secreted, (semaphorin) 3A | 0.79 | 0.004 |
| *Gbp5* | guanylate binding protein 5 | 0.79 | 0.015 |
| *Cckbr* | cholecystokinin B receptor | 0.78 | 0.004 |
| *Slc6a20a* | solute carrier family 6 (neurotransmitter transporter), member 20A | 0.77 | 0.004 |
| *Lrrc32* | leucine rich repeat containing 32 | 0.76 | 0.026 |
| *Tnnt2* | troponin T2, cardiac | 0.76 | 0.024 |
| *Irgm2* | immunity-related GTPase family M member 2 | 0.76 | 0.013 |
| *Mrc2* | mannose receptor, C type 2 | 0.75 | 0.004 |
| *Blnk* | B cell linker | 0.74 | 0.022 |
| *Igfn1* | immunoglobulin-like and fibronectin type III domain containing 1 | 0.74 | 0.004 |
| *A430107O13Rik* | cadherin-like and PC-esterase domain containing 1 | 0.74 | 0.015 |
| *Vwf* | Von Willebrand factor | 0.73 | 0.004 |
| *Ifitm3* | interferon induced transmembrane protein 3 | 0.73 | 0.004 |
| *Cfh* | complement component factor h | 0.72 | 0.004 |
| *Cobl* | cordon-bleu WH2 repeat | 0.72 | 0.004 |
| *Adra1b* | adrenergic receptor, alpha 1b | 0.72 | 0.037 |
| *Syt2* | synaptotagmin II | 0.71 | 0.004 |
| *H2-K1* | histocompatibility 2, K1, K region | 0.71 | 0.004 |
| *Thbd* | thrombomodulin | 0.70 | 0.004 |
| *Slc22a8* | solute carrier family 22 (organic anion transporter), member 8 | 0.69 | 0.004 |
| *E130203B14Rik* | V-set and transmembrane domain containing 4 | 0.69 | 0.042 |
| *Sowahb* | sosondowah ankyrin repeat domain family member B | 0.69 | 0.029 |
| *Lyz2* | lysozyme 2 | 0.68 | 0.004 |
| *Rgs6* | regulator of G-protein signaling 6 | 0.68 | 0.018 |
| *Cyp1b1* | cytochrome P450, family 1, subfamily b, polypeptide 1 | 0.68 | 0.033 |
| *Colec12* | collectin sub-family member 12 | 0.67 | 0.004 |
| *Slc17a6* | solute carrier family 17 (sodium-dependent inorganic phosphate cotransporter), member 6 | 0.67 | 0.004 |
| *Ntsr1* | neurotensin receptor 1 | 0.66 | 0.029 |
| *Vill* | villin-like | 0.66 | 0.020 |
| *Myoc* | myocilin | 0.66 | 0.004 |
| *Rims3* | regulating synaptic membrane exocytosis 3 | 0.66 | 0.010 |
| *Sv2c* | synaptic vesicle glycoprotein 2c | 0.65 | 0.022 |
| *Coch* | cochlin | 0.64 | 0.010 |
| *Bmp6* | bone morphogenetic protein 6 | 0.64 | 0.013 |
| *H2-D1* | histocompatibility 2, D region locus 1 | 0.63 | 0.004 |
| *Kdm6a* | lysine (K)-specific demethylase 6A | 0.63 | 0.004 |
| *Nid1* | nidogen 1 | 0.63 | 0.007 |
| *Serpinf1* | serine (or cysteine) peptidase inhibitor, clade F, member 1 | 0.62 | 0.015 |
| *Tmem90a* | synapse differentiation inducing 1 like | 0.62 | 0.031 |
| *Ranbp3l* | RAN binding protein 3-like | 0.62 | 0.018 |
| *Plxdc1* | plexin domain containing 1 | 0.61 | 0.004 |
| *Gsg1l* | GSG1-like | 0.60 | 0.004 |
| **Genes higher in males at baseline** | | | |
| **Gene** | **Gene Name** | **Log2 Fold Change** | **P-value** |
| *Fibcd1* | fibrinogen C domain containing 1 | -0.59 | 0.004 |
| *Npr3* | natriuretic peptide receptor 3 | -0.59 | 0.004 |
| *Calb2* | calbindin 2 | -0.59 | 0.004 |
| *Mest* | mesoderm specific transcript | -0.59 | 0.004 |
| *Slc9a4* | solute carrier family 9 (sodium/hydrogen exchanger), member 4 | -0.60 | 0.010 |
| *Crym* | crystallin, mu | -0.61 | 0.004 |
| *Nfkbiz* | nuclear factor of kappa light polypeptide gene enhancer in B cells inhibitor, zeta | -0.61 | 0.018 |
| *Irs2* | insulin receptor substrate 2 | -0.61 | 0.004 |
| *Cpne2* | copine II | -0.62 | 0.004 |
| *Amigo2* | adhesion molecule with Ig like domain 2 | -0.63 | 0.004 |
| *Tob1* | transducer of ErbB-2.1 | -0.66 | 0.004 |
| *Fbxo33* | F-box protein 33 | -0.66 | 0.004 |
| *Spty2d1* | SPT2, Suppressor of Ty, domain containing 1 (S. cerevisiae) | -0.68 | 0.004 |
| *Cpne7* | copine VII | -0.68 | 0.004 |
| *Nfil3* | nuclear factor, interleukin 3, regulated | -0.69 | 0.007 |
| *Jun* | jun proto-oncogene | -0.69 | 0.004 |
| *Sgk1* | serum/glucocorticoid regulated kinase 1 | -0.70 | 0.004 |
| *Hif3a* | hypoxia inducible factor 3, alpha subunit | -0.70 | 0.004 |
| *Gm5177* | glyceraldehyde-3-phosphate dehydrogenase pseudogene | -0.71 | 0.004 |
| *Dlk1* | delta-like 1 homolog (Drosophila) | -0.72 | 0.040 |
| *Enpp2* | ectonucleotide pyrophosphatase/phosphodiesterase 2 | -0.73 | 0.004 |
| *Zcchc5* | zinc finger, CCHC domain containing 5 | -0.74 | 0.031 |
| *Gm129* | circadian associated repressor of transcription | -0.74 | 0.004 |
| *Arl5b* | ADP-ribosylation factor-like 5B | -0.74 | 0.004 |
| *Nr4a3* | nuclear receptor subfamily 4, group A, member 3 | -0.75 | 0.004 |
| *Scml4* | sex comb on midleg-like 4 (Drosophila) | -0.76 | 0.040 |
| *Trhr* | thyrotropin releasing hormone receptor | -0.81 | 0.010 |
| *Mc4r* | melanocortin 4 receptor | -0.82 | 0.022 |
| *Rasd1* | RAS, dexamethasone-induced 1 | -0.82 | 0.026 |
| *Per1* | period circadian clock 1 | -0.83 | 0.004 |
| *Adamts14* | a disintegrin-like and metallopeptidase (reprolysin type) with thrombospondin type 1 motif, 14 | -0.83 | 0.046 |
| *Csrnp1* | cysteine-serine-rich nuclear protein 1 | -0.85 | 0.004 |
| *Ppp1r15a* | protein phosphatase 1, regulatory (inhibitor) subunit 15A | -0.86 | 0.004 |
| *Col8a1* | collagen, type VIII, alpha 1 | -0.91 | 0.015 |
| *Mid1* | midline 1 | -0.91 | 0.004 |
| *Kl* | klotho | -0.91 | 0.004 |
| *Glp2r* | glucagon-like peptide 2 receptor | -0.93 | 0.004 |
| *Dio3* | deiodinase, iodothyronine type III | -0.96 | 0.024 |
| *Gadd45b* | growth arrest and DNA-damage-inducible 45 beta | -0.98 | 0.004 |
| *Otx2* | orthodenticle homeobox 2 | -1.00 | 0.018 |
| *Dusp5* | dual specificity phosphatase 5 | -1.01 | 0.004 |
| *Rgs2* | regulator of G-protein signaling 2 | -1.02 | 0.004 |
| *Clic6* | chloride intracellular channel 6 | -1.08 | 0.004 |
| *Gpr101* | G protein-coupled receptor 101 | -1.09 | 0.004 |
| *Dusp6* | dual specificity phosphatase 6 | -1.10 | 0.004 |
| *Sostdc1* | sclerostin domain containing 1 | -1.10 | 0.004 |
| *Zfp36* | zinc finger protein 36 | -1.18 | 0.004 |
| *Nr4a2* | nuclear receptor subfamily 4, group A, member 2 | -1.20 | 0.004 |
| *Pcdh8* | protocadherin 8 | -1.22 | 0.004 |
| *Apold1* | apolipoprotein L domain containing 1 | -1.23 | 0.004 |
| *Serpine1* | serine (or cysteine) peptidase inhibitor, clade E, member 1 | -1.23 | 0.026 |
| *Adamts1* | a disintegrin-like and metallopeptidase (reprolysin type) with thrombospondin type 1 motif, 1 | -1.23 | 0.004 |
| *Maff* | v-maf musculoaponeurotic fibrosarcoma oncogene family, protein F (avian) | -1.26 | 0.007 |
| *Tiparp* | TCDD-inducible poly(ADP-ribose) polymerase | -1.27 | 0.004 |
| *Rasl11a* | RAS-like, family 11, member A | -1.33 | 0.004 |
| *Krt18* | keratin 18 | -1.35 | 0.010 |
| *Kcnj2* | potassium inwardly-rectifying channel, subfamily J, member 2 | -1.35 | 0.004 |
| *Gm13889* | predicted gene 13889 | -1.43 | 0.004 |
| *Fosl2* | fos-like antigen 2 | -1.44 | 0.004 |
| *Egr3* | early growth response 3 | -1.48 | 0.004 |
| *Chrm5* | cholinergic receptor, muscarinic 5 | -1.49 | 0.018 |
| *Slc39a4* | solute carrier family 39 (zinc transporter), member 4 | -1.49 | 0.004 |
| *Sik1* | salt inducible kinase 1 | -1.56 | 0.004 |
| *1500015O10Rik* | RIKEN cDNA 1500015O10 gene | -1.66 | 0.004 |
| *Steap1* | six transmembrane epithelial antigen of the prostate 1 | -1.66 | 0.013 |
| *Errfi1* | ERBB receptor feedback inhibitor 1 | -1.67 | 0.004 |
| *Ier2* | immediate early response 2 | -1.73 | 0.004 |
| *Dusp1* | dual specificity phosphatase 1 | -1.73 | 0.004 |
| *Atf3* | activating transcription factor 3 | -1.74 | 0.004 |
| *Vgll3* | vestigial like family member 3 | -1.75 | 0.004 |
| *Ptgs2* | prostaglandin-endoperoxide synthase 2 | -1.85 | 0.004 |
| *F5* | coagulation factor V | -1.93 | 0.004 |
| *Cldn2* | claudin 2 | -1.99 | 0.004 |
| *Gadd45g* | growth arrest and DNA-damage-inducible 45 gamma | -2.04 | 0.004 |
| *Aqp1* | aquaporin 1 | -2.06 | 0.004 |
| *Folr1* | folate receptor 1 (adult) | -2.11 | 0.004 |
| *Egr1* | early growth response 1 | -2.12 | 0.004 |
| *Nr4a1* | nuclear receptor subfamily 4, group A, member 1 | -2.24 | 0.004 |
| *Gabra6* | gamma-aminobutyric acid (GABA) A receptor, subunit alpha 6 | -2.29 | 0.004 |
| *Cyr61* | cysteine rich protein 61 | -2.42 | 0.004 |
| *Arc* | activity regulated cytoskeletal-associated protein | -2.44 | 0.004 |
| *Arl4d* | ADP-ribosylation factor-like 4D | -2.46 | 0.004 |
| *Junb* | jun B proto-oncogene | -2.48 | 0.004 |
| *Kcne2* | potassium voltage-gated channel, Isk-related subfamily, gene 2 | -2.52 | 0.004 |
| *Slc4a5* | solute carrier family 4, sodium bicarbonate cotransporter, member 5 | -2.62 | 0.004 |
| *Egr4* | early growth response 4 | -2.85 | 0.004 |
| *Ttr* | transthyretin | -2.89 | 0.004 |
| *Tmem72* | transmembrane protein 72 | -3.17 | 0.004 |
| *Btg2* | B cell translocation gene 2, anti-proliferative | -3.47 | 0.004 |
| *Npas4* | neuronal PAS domain protein 4 | -4.36 | 0.004 |
| *Fosb* | FBJ osteosarcoma oncogene B | -4.70 | 0.004 |
| *Fos* | FBJ osteosarcoma oncogene | -5.57 | 0.004 |
| *Egr2* | early growth response 2 | -6.18 | 0.004 |
| *Eif2s3y* | eukaryotic translation initiation factor 2, subunit 3, structural gene Y-linked | -8.31 | 0.004 |
| *Ddx3y* | DEAD (Asp-Glu-Ala-Asp) box polypeptide 3, Y-linked | -9.58 | 0.040 |
