## Supplementary Table 7. Long-term_Male and Female Biased Genes. for "Enduring and sex-specific changes in hippocampal gene expression after a subchronic immune challenge"

**Supplementary Table 7. Male-biased and female-biased genes that are differentially expressed three months after subchronic immune challenge (Long-term condition).**

| ***MALE-BIASED GENES*** | | |
| --- | --- | --- |
| **UPREGULATED** | **Males** | **Females** |
|  | *Cldn2* | *Dlk1* |
|  | *Clic6* | *Fos* |
|  | *Col8a1* | *Gpr101* |
|  | *Enpp2* | *Mid1* |
|  | *Folr* | *NPAS4* |
|  | *Kcne2* |  |
|  | *Krt18* |  |
|  | *Slc39a4* |  |
|  | *Slc4a5* |  |
|  | *Steap1* |  |
|  | *Tmem72* |  |
|  | *Ttr* |  |
| **DOWN-**  **REGULATED** | *Csrnp1* | **N/A** |
|  | *Arl5b* |  |
|  | *Nfkbiz* |  |
|  | *Per1* |  |
|  | *Gadd45b* |  |
|  | *Dusp6* |  |
|  | *Ppp1r15a* |  |
|  | *Nfil3* |  |
|  | *Adamts1* |  |
|  | *Tiparp* |  |
|  | *Rasl11a* |  |
|  | *Pcdh3* |  |
|  | *Fosl2* |  |
|  | *Sik1* |  |
|  | *Apold1* |  |
|  | *Nr4a2* |  |
|  | *Egr3* |  |
|  | *Dusp1* |  |
|  | *Ptgs2* |  |
|  | *Gadd45g* |  |
|  | *Ier2* |  |
|  | *Atf3* |  |
|  | *Egr1* |  |
|  | *Nr4a1* |  |
|  | *Arc* |  |
|  | *Junb* |  |
|  | *Cyr61* |  |
|  | *Arl4d* |  |
|  | *Egr4* |  |
|  | *Btg2* |  |
|  | *Npas4* |  |
|  | *Fos* |  |
|  | Fosb |  |
|  | Egr2 |  |
| ***FEMALE-BIASED GENES*** | | |
| **UPREGULATED** | **Males** | **Females** |
|  | *Aebp1* | **N/A** |
|  | *Aldh1a2* |  |
|  | *Bmp6* |  |
|  | *Bmp7* |  |
|  | *C2* |  |
|  | *Carbp2* |  |
|  | *Cd74* |  |
|  | *Coch* |  |
|  | *Col1a1* |  |
|  | *Col1a2* |  |
|  | *Col3a1* |  |
|  | *Col4a6* |  |
|  | *Col9a2* |  |
|  | *Efemp1* |  |
|  | *Emilin1* |  |
|  | *Fgfbp1* |  |
|  | *Fmod* |  |
|  | *Gjb2* |  |
|  | *Gpr81* |  |
|  | *H2-Ab1* |  |
|  | *Ifit1* |  |
|  | *Ifitm3* |  |
|  | *Islr* |  |
|  | *Itih2* |  |
|  | *Mpzl2* |  |
|  | *Mrc1* |  |
|  | *Mrc2* |  |
|  | *Oasl2* |  |
|  | *Ogn* |  |
|  | *Prdm6* |  |
|  | *Ptgdr* |  |
|  | *Ptgds* |  |
|  | *Serping1* |  |
|  | *Slc13a3* |  |
|  | *Slc22a6* |  |
|  | *Slc47a1* |  |
|  | *Slc6a12* |  |
|  | *Slc6a13* |  |
|  | *Slc6a20a* |  |
|  | *Sned1* |  |
|  | *Sphk1* |  |
| **DOWN-**  **REGULATED** | Cnp11 | *Adora2a* |
|  |  | *Adra1b* |
|  |  | *Cckbr* |
|  |  | *Coch* |
|  |  | *Drd2* |
|  |  | *Foxp2* |
|  |  | *GBP4* |
|  |  | *Gpr88* |
|  |  | *ifit1* |
|  |  | *Irgm2* |
|  |  | *rasgef1c* |
|  |  | *Scn4b* |
|  |  | *Spp1* |
