## Supplementary Table 8. Long-term + Acute_Male and Female Biased Genes. for "Enduring and sex-specific changes in hippocampal gene expression after a subchronic immune challenge"

**Supplementary Table 8. Male-biased and female-biased genes that are differentially expressed in response to an acute immune challenge, three months after prior subchronic immune challenge (Long-term + Acute condition).**

| ***MALE-BIASED GENES*** | | |
| --- | --- | --- |
|  | **Males** | **Females** |
|  | Gene | Gene |
| **UPREGULATED** | *Dlk1* | **N/A** |
|  | *Sgk1* |  |
| **DOWN-REGULATED** | **N/A** | Cldn2 |
|  |  | Col8a1 |
|  |  | F5 |
| ***FEMALE-BIASED GENES*** | | |
|  | **Males** | **Females** |
|  | Gene | Gene |
| **UPREGULATED** | *Adora2a* | *Blnk* |
|  | *Col6a4* | *Cckbr* |
|  | *Drd1a* | *Cd4* |
|  | *Drd2* | *Cobl* |
|  | *Foxp2* | *Dpp4* |
|  | *Igfn1* | *Drd1a* |
|  | *Sphk1* | *Drd2* |
|  |  | *Foxp2* |
|  |  | *Gpr88* |
|  |  | *Igfbp6* |
|  |  | *Plxdc1* |
|  |  | *Rasgef1c* |
|  |  | *Rgs6* |
|  |  | *Rims3* |
|  |  | *Scn4b* |
|  |  | *Slc6a12* |
|  |  | *Sowahb* |
|  |  | *Syt2* |
|  |  | *Tmem90a* |
|  |  | *Trhr2* |
|  |  | *Wnt10a* |
|  | **Males** | **Females** |
|  | Gene | Gene |
| **DOWN-REGULATED** | *Islr* | *Aldh1a2* |
|  |  | *Aox3* |
|  |  | *Bmp6* |
|  |  | *Bmp7* |
|  |  | *Cd74* |
|  |  | *Cdh1* |
|  |  | *Cldn2* |
|  |  | *Clic6* |
|  |  | *Col8a1* |
|  |  | *Cpne7* |
|  |  | *Crym* |
|  |  | *Cyp1b1* |
|  |  | *Dlk1* |
|  |  | *Enpp2* |
|  |  | *F5* |
|  |  | *Fibcd1* |
|  |  | *Fmod* |
|  |  | *Folr1* |
|  |  | *Gbp2* |
|  |  | *Gjb2* |
|  |  | *Gpr101* |
|  |  | *H2-Aa* |
|  |  | *H2-Ab1* |
|  |  | *H2-Q1* |
|  |  | *Hif3a* |
|  |  | *Ifi44* |
|  |  | *Ifitm3* |
|  |  | *Itih2* |
|  |  | *Kcne2* |
|  |  | *Kl* |
|  |  | *Krt18* |
|  |  | *Mc4r* |
|  |  | *Mid1* |
|  |  | *Mpzl2* |
|  |  | *Mrc1* |
|  |  | *Npr3* |
|  |  | *Oasl2* |
|  |  | *Otx2* |
|  |  | *Pcolce* |
|  |  | *Ptgds* |
|  |  | *Serping1* |
|  |  | *Slc13a4* |
|  |  | *Slc39a4* |
|  |  | *Slc4a5* |
|  |  | *Slc6a13* |
|  |  | *Slc6a20a* |
|  |  | *Slc9a4* |
|  |  | *Sostdc1* |
|  |  | *Spp1* |
|  |  | *Steap1* |
|  |  | *Tmem72* |
|  |  | *Ttr* |
